# The Insertion of an ATTTC Repeat in an Alu Element Hyperactivates a Primate-Specific Neurodevelopmental Enhancer in Spinocerebellar Ataxia Type 37

**DOI:** 10.1101/2025.06.03.657633

**Authors:** Joana R. Loureiro, Ana F. Castro, Ana Sofia Figueiredo, Ana Eufrásio, Ashutosh Dhingra, Mafalda Galhardo, Hugo Marcelino, Catarina Rodrigues, Paula Sampaio, Maria Azevedo, Mafalda Sousa, Sofia Dória, Patrizia Rizzu, Peter Heutink, José Bessa, Isabel Silveira

**Affiliations:** Genetics of Cognitive Dysfunction Laboratory, i3S-Instituto de Investigação e Inovação em Saúde, Universidade do Porto, 4200-135 Porto, Portugal; IBMC-Institute for Molecular and Cell Biology, Universidade do Porto, 4200-135 Porto, Portugal; ICBAS, Universidade do Porto, 4050-313 Porto, Portugal; Vertebrate Development and Regeneration Laboratory, i3S-Instituto de Investigação e Inovação em Saúde, Universidade do Porto, 4200-135 Porto, Portugal; German Center for Neurodegenerative Diseases, 72076 Tübingen, Germany; Advanced Light Microscopy, i3S-Instituto de Investigação e Inovação em Saúde, Universidade do Porto, 4200-135 Porto, Portugal; Genetic Unit, Department of Pathology, Faculty of Medicine and RISE-Health, University of Porto, Porto, Portugal

**Keywords:** Pentanucleotide Repeat expansion, Short Tandem Repeat (STR), Spinocerebellar Ataxia Type 37 (SCA37), Neurodegenerative Disease, *DAB1* Primate-specific Alu Element, Axonal Pathfinding, Cerebellar Transcriptional Enhancer, iPSN, Circularized Chromosome Conformation Capture Sequencing (4C-seq), Cap Analysis of Gene Expression (CAGE)

## Abstract

Alu are evolutionarily very old primate-specific interspersed repeat elements that constitute ∼11% of the human genome. They are a source of short tandem repeats (STRs), which often expand in size and originate inherited neuromuscular and neurodegenerative disorders. How expanded STR insertion mutations within Alu STRs culminate in disease remains unknown. Here we report an Alu STR located in an intron of *DAB1* that functions as a neurodevelopmental enhancer. We demonstrated that an ATTTC repeat insertion in this *DAB1* Alu STR, known to cause spinocerebellar ataxia type 37 (SCA37), hyperactivates a neurodevelopmental *DAB1* enhancer. Importantly, we showed that neurons derived from SCA37 subjects have higher levels of *DAB1* expression and DAB1 overexpression causes abnormal axonal pathfinding *in vivo*. Overall, these results establish that neuronal dysregulation of a developmental *DAB1* Alu STR enhancer contributes to SCA37 pathogenesis, an unexplored mechanism likely acting in many Alu STR diseases, potentially reshaping the therapeutic landscape.

## Introduction

Alu elements are retrotransposons with ∼1.2 million copies in the human genome, belonging to the type of primate-specific short interspersed nucleotide elements^1^. They are composed of two monomers, left and right, separated by a polymorphic middle A-rich region and an ending poly-A tail tract of variable size. These polymorphic sequences are a cradle of short tandem repeats (STRs), or microsatellites, which are 1-6 base pair motifs of repeated units of DNA^1,2^. The Alu left monomers have an internal RNA polymerase III promoter that when not degenerated drives Alu transcription^3^. The evolutionary older transcriptionally expressed elements of this family of retrotransposons could have been recruited by the host genome to function as transcriptional enhancers for nearby protein-coding genes^4^.

STR expansions and insertions cause over 60 neurodevelopmental, neurodegenerative and/or neuromuscular disorders, including 17 types of spinocerebellar ataxia (SCA)^5,6^. At least 13 expansions and insertions causing disorders have arisen from Alu STRs^6,7^. We have discovered one of these expanded insertions in an Alu STR as the genetic basis of SCA37^8^. An (ATTTC)_n_ was found within a preexisting (ATTTT)_n,_ in the middle A-rich tract belonging to an old degenerated antisense AluJb, located in the intronic region of *DAB1*. For the non-pathogenic alleles, we have seen a simple [(ATTTT)_7-400_] sequence, whereas pathogenic expanded alleles presented a complex [(ATTTT)_60-79_(ATTTC)_31-75_(ATTTT)_58-90_] configuration^8^ observed also in most affected subjects from other studies^9,10^. This repeat can, however, expand to much larger pathogenic alleles that greatly complicate their detection^11,12^. In fact, with advances in genomics and bioinformatics analysis, (ATTTC)_n_ insertions have been found in distinct genes causing at least seven additional pathologies classified as familial adult myoclonic epilepsies^12–17^.

*DAB1* encodes a reelin adaptor protein, that is an obligate effector of the Reelin signaling pathway, crucial for neuronal migration and cerebellar cell layer organization during development^18,19^. A developmental delay in dendrite outgrowth and axonal branching has been seen in *Dab1* mutant mice with mispositioning of neurons in the cerebellum, hippocampus and cortex, and presenting a motor phenotype of ataxia^18–20^. Dab1 is highly enriched in axonal growth cones during pathfinding, in development, contributing to the final neuronal positioning in the mammalian brain^21^. Later on, in the young adult brain, Dab1 continues to regulate the maturation of dendritic spines, synaptogenesis and astroglial ensheathment of synapses^22^.

The human *DAB1* gene contains several alternative first exons, which result in transcripts with variable UTRs encompassing the (ATTTC)_n_ region^8^. In *DAB1*, the (ATTTC)_n_ insertion is located in intronic regions of transcripts with cerebellar-specific expression. Their expression starts early during fetal brain development and is maintained in the adult cerebellum^8^. In SCA37 cerebellar autopsy tissue, *DAB1* transcripts have shown increased expression compared with tissue from age-matched elderly controls^9^. Notwithstanding the observed overexpression of DAB1 protein in cerebellar tissue from SCA37 ageing subjects^9^, whether this plays a role in early pathogenesis remains unknown. This disease shows a variable age of onset, ranging from the second to the sixtieth decade of life, and heterogeneous clinical presentation, usually of progressive gait, limb, and speech ataxia, seldom associated with dementia or motor neuron pathology^8,9,11^. This variability in onset is explained by the unstable nature of this type of pathogenic repeat mutations. They trigger earlier, sometimes congenital, and more severe symptoms when the pathogenic repeat size is larger^8^, which is a classical hallmark of repeat expansion conditions^23–26^. (ATTTC)_n_ insertions in Alu STRs are also the molecular basis of at least six more familial neurological disorders^7,12,27^. How the pathological pentanucleotide repeat acts to trigger these pathologies remains to be elucidated.

Here we report that the intronic (ATTTC)_n_ insertion in an Alu STR, within the *DAB1* repeat locus (DRL), hyperactivates a neurodevelopmental enhancer in SCA37. We demonstrated that the DRL interacts with cerebellar-specific *DAB1* promoters and enhances *DAB1* transcriptional expression, which is upregulated in SCA37-derived neurons. DAB1 overexpression during zebrafish development causes abnormal axonal pathfinding. These results highlight the crucial role of Alu STRs in regulating gene expression and cellular function, extending its relevance further to other diseases.

## Methods

### Experimental Model and Subject Details

#### Cell line generation and cell culture

Primary cultures of fibroblasts established from skin biopsies as previously described^8^ and human embryonic kidney 293T (HEK293T) cell line (ATCC #CRL-3216) were cultured in DMEM GlutaMAX medium (#31966047, Gibco, Thermo Fisher Scientific) with 10% fetal bovine serum (#A5256801, Gibco, Thermo Fisher Scientific) and 100 units/mL penicillin, and 0.1 mg/mL streptomycin (#15140122, Gibco, Thermo Fisher Scientific), at 37°C, with 5% CO_2_. SCA37 induced pluripotent stem cells (iPSC) were reprogrammed from fibroblasts of three affected individuals with the ATTTC repeat mutation, by an integration-free protocol overexpressing OCT3/4, SOX2, KLF4, L-MYC and LIN28, as described^28^. The generated SCA37 iPSC clones (Table S1) were karyotyped and pluripotency was confirmed (detailed in “Immunostaining”). Control iPSC ND41866 and GM23280 were obtained from Coriell institute, NAS6 was a gift from Dr T. Kunath, University of Edinburgh^29^ (Table S1). iPSC lines were maintained in Essential 8 Flex Medium kit (#A2858501; Gibco, Thermo Fisher Scientific), in plates precoated with Corning Matrigel hESC-Qualified Matrix (#354277; Corning). SCA37 and control iPSC lines were differentiated into cerebral cortical glutamatergic neurons, by neurogenin 2 (NGN2) overexpression, as previously described^30,31^, until day 8. The expression of protein DAB1 and the neuronal markers TUJ1 and MAP2 was confirmed by immunofluorescence (detailed in “Immunostaining”). This project was approved by the i3S ethics committee (N3/CECRI/2020).

Human neural stem cells (hNSC; line CB192)^32^ were maintained in DMEM-F12 GlutaMAX (#31331028, Gibco, Thermo Fisher Scientific) supplemented with 1x N-2 (#17502048; Gibco, Thermo Fisher Scientific), 0.05x B27 (#17504044, Gibco, Thermo Fisher Scientific), 10 ng/mL EGF (#315-09; Peprotech, Thermo Fisher Scientific); 10 ng/mL FGF (# 100-18B, Peprotech, Thermo Fisher Scientific); 1µg/mL Laminin (#L2020-1MG; Sigma-Aldrich, Merck) and 100 units/mL penicillin and 0.1 mg/mL streptomycin (#15140122; Gibco, Thermo Fisher Scientific) in T-flasks or plates pre-coated with 0.01% Poly-L-Lysin (#P8920-100ML, Sigma-Aldrich, Merck). DAB1 protein expression was also confirmed by immunofluorescence (detailed in “Immunostaining”). The cultured cells were regularly tested for the absence of mycoplasma contamination, by PCR.

#### Zebrafish animals and husbandry

Zebrafish (*Danio rerio*) experiments complied with standard animal care guidelines and national legislation for animal experimentation (Decreto-Lei n° 113/2013) and the European Union legislation (Directive 2010/63/EU), being authorized by the i3S animal ethics committee (ORBEA - Órgãos Responsáveis pelo Bem-Estar dos Animais, ref. 2018-29) and Direção-Geral de Alimentação e Veterinária (DGAV; Ref. 022872/2020-12-31). Maintenance and handling of zebrafish animals were performed in the i3S zebrafish animal facility (licensed by DGAV and part of the AAALAC - International accredited animal care and use program). Wild-type zebrafish with Oregon AB genetic background were crossed for the generation of embryos for microinjection procedures, except for axonal analysis in larvae cerebella, which was performed in gSA2AzGFF152B; UAS:GFP transgenic line^33,34^.

## Method Details

### iPSC karyotype analysis

iPSCs karyotype analysis was performed to assess chromosomal stability and detect any abnormalities, as reported^35^. Cells were maintained in log-phase growth and harvested at approximately 70% confluency for karyotype analysis. To induce metaphase arrest, iPSCs were treated with 10 µg/mL KaryoMAX™ Colcemid™ solution (#15210040, Gibco, Thermo Fisher Scientific) for 1 hour at 37°C. Following treatment, iPSCs were dissociated into a single-cell suspension using TrypLE (#12605-028, Gibco, Thermo Fisher Scientific), transferred to a tube containing fetal bovine serum (#A5256801, Gibco, Thermo Fisher Scientific), and centrifuged at 150 × g for 8 min. The supernatant was removed, and cells were gently resuspended in 0.075 M KCl (#10575090, Thermo Fisher Scientific) dropwise and incubated at 37°C for 25 min. For fixation, the cell suspension was treated with pre-cooled fixative solution (3:1 methanol (#1.06009.1000, Merck) to glacial acetic acid 100% (#1.00063.2511, Merck)) for 10 min, at room temperature, followed by centrifugation at 150 × g for 8 min. The supernatant was carefully removed, and the fixation step was repeated with an incubation of 30 min, at room temperature. After the final centrifugation, the pellet was resuspended in 1.5 mL of fixative solution and stored at 4°C. Preparation of chromosome slides and GTL-banding using Leishman stain (#L6254, Sigma-Aldrich, Merck) were performed according to standard methods. At least 20 metaphases were counted for each iPSC line, and the International System for Human Cytogenomic Nomenclature (ISCN) rules were followed^36^. Metaphases were karyotyped using the CytoVision/v3.932 software (Applied Imaging Cytovision).

### Zebrafish transgenesis and microinjection

Zebrafish transgeneses were performed according to the *Tol2* transposon system^37^ by co-injection of 5 nL of each test vector with Tol2 mRNA (25 ng each) and 10% phenol red (#P0290, Sigma-Aldrich, Merck) in one-to two-cell stage zebrafish embryos. The consequences of DAB1 overexpression in zebrafish were assessed by injection of *in vitro* synthesized *DAB1* mRNA (detailed in “*In vitro* RNA synthesis”), at 100 ng/µL, in wild-type embryos, and at 200 ng/µL in gSA2AzGFF152B; UAS:GFP transgenic embryos. Embryos were cultured at 28°C in Petri dishes containing E3 medium (5 mM NaCl (#MB15901, NZYtech), 0.17 mM KCl (#P9541, Sigma-Aldrich, Merck), 0.33 mM CaCl2 (#C3881, Sigma-Aldrich, Merck), 0.33 mM MgSO4 (#63140, Sigma-Aldrich, Merck), 0.00015% methylene blue (#A1402, PanReac AppliChem ITW Reagents)) supplemented with 0.003% 1-phenyl-2-thiourea (#P7629, Sigma-Aldrich, Merck) to delay pigmentation formation^1^. Embryos and larvae were anaesthetized with 150 mg/L of tricaine (ethyl 3-aminobenzoate; #E10521, Sigma-Aldrich, Merck) added to the E3 medium and the phenotypes were documented.

### *In Vitro* RNA Synthesis

Transposase *Tol2* RNA was *in vitro* synthesized from pCSF2A-Tol2 plasmid with SP6 RNA polymerase (#EP0131, Thermo Fisher Scientific). Human *DAB1* mRNA was *in vitro* transcribed from pCMV6-XL4-DAB1 mammalian expression vector (#SC113027, Origene) with T7 RNA polymerase (#EP0111, Thermo Fisher Scientific). Before *in vitro* RNA synthesis of *Tol2* and *DAB1*, plasmid DNA was linearized with NotI (#IVGN0016, Anza, Thermo Fisher Scientific) or XhoI (#IVGN0086, Anza, Thermo Fisher Scientific), respectively, and purified with DNA Clean & Concentrator kit (#D4013, ZYMO Research). RNAs were purified with RNA Clean & Concentrator kit (#R1017, ZYMO Research) prior to zebrafish injection.

### Circularized Chromosome Conformation Capture sequencing (4C-seq)

4C-seq was performed in iPSC lines derived from two affected individuals (AF2Cl4 and AF4Cl32) and two unaffected controls (ND41866 and GM23280), as previously described^38^ with few modifications. The restriction enzymes DpnII (#R0543M, NEB) and Csp6I (#ER0211, Thermo Fisher Scientific) were used in the first and second restriction, respectively. Circularized chromatin was purified by Amicon Ultra-15 Centrifugal Filter Device (#UFC901024, Milipore, Merck). 4C-seq PCR was performed with Expand Long Template Polymerase (#11759060001, Roche, Merck) with primers containing Illumina adapters and indexes suitable for pair-end sequencing (Table S2). 4C-seq PCR products were purified with the High Pure PCR Product Purification Kit (#11796828001, Roche, Merck) followed by AMPure XP PCR purification kit (#B37419AB, Agencourt AMPure XP). 4C-seq libraries were sequenced in Illumina HiSeqX; the processing and alignment were performed against the human genome hg38, as previously described^39,40^. More than 6 million reads per sample aligned to the human genome (hg38, Fig.1); or >1 million reads aligned, Supp.Fig.2) with Bowtie2^41^ in pair end mode with parameters "-X 2000 -- very-sensitive" (global alignment mode). Valid reads within fragments >40 bp flanked by cutting sites of the restriction enzymes (DpnII - GATC and Csp6I - GTAC) were counted and smoothed using the average over 30 bp units to decrease signal sparsity. Genomic regions with signal above expected based on a theoretical Poisson distribution were defined as targets representing regions that interact with the viewpoint.

**Figure 1.**
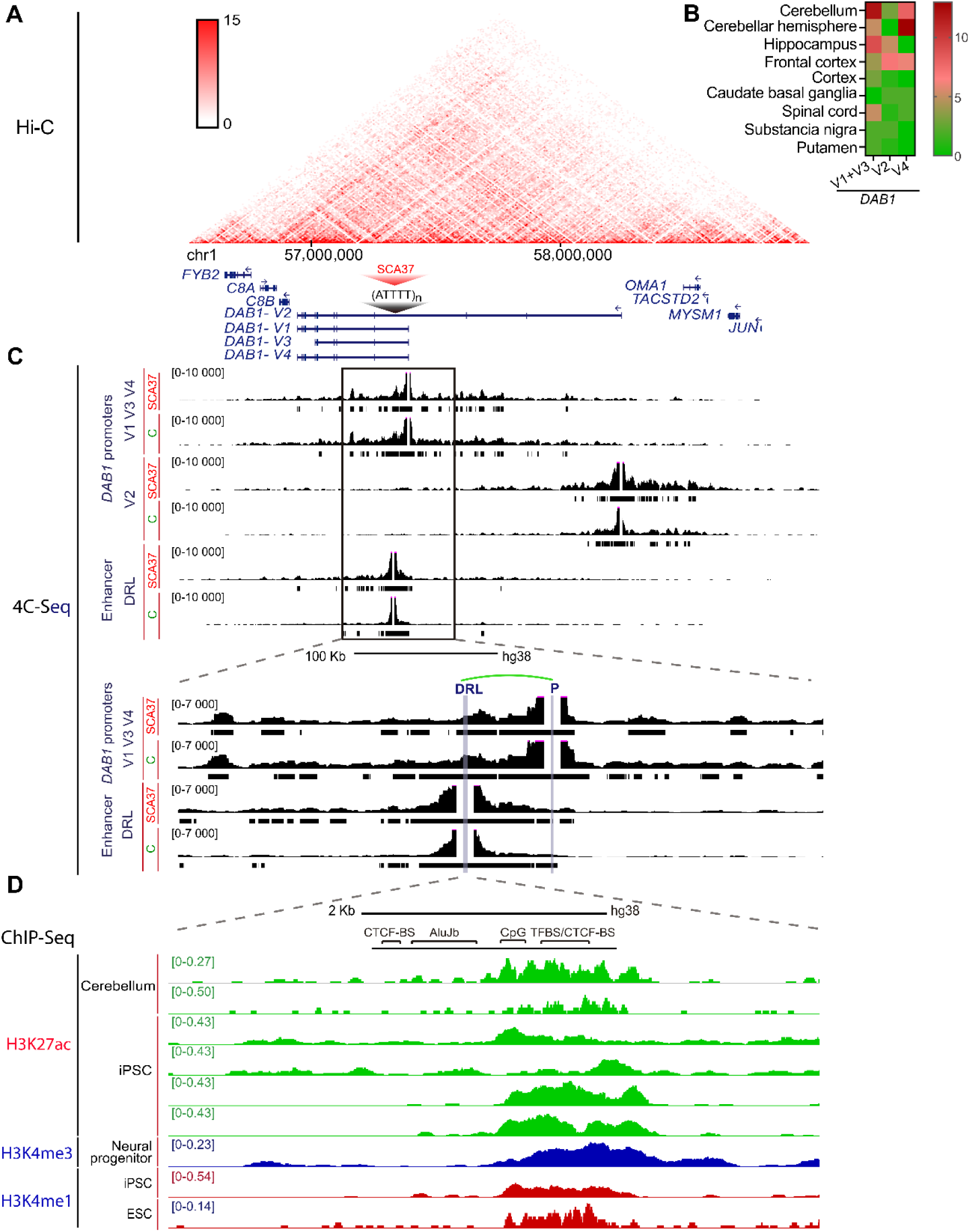
Regulatory landscape of the *DAB1* gene and enhancer marks at the DRL region. (A) Hi-C data from human cerebellar cells, obtained from the ENCODE project, visualized with the 3D Genome Browser, at 10 kb resolution showing the TAD organization of *DAB1* (hg38, chr1:56,800,000-58,700,000); *DAB1* mRNA variants and (ATTTT)_n_ location are depicted. (B) Expression levels of *DAB1* mRNA variants across different brain regions obtained from GTEx. (C) 4C-seq interaction profiles of representative iPSCs derived from SCA37 affected and unaffected individuals with viewpoints in the promoters of *DAB1* V1, V3 and V4 transcripts and that of V2 *DAB1* variant, as well as the DRL region; for each sample is shown the normalized read coverage with statistically significant interactions below. (D) Human cells and tissues with DRL positive ChIP-seq signal the for H3K27ac, H3K4me3 and H3K4me1, from ChIP-Atlas. C-control, P-promoter. See also Figures S1, S2 and S3.

**Figure 2.**
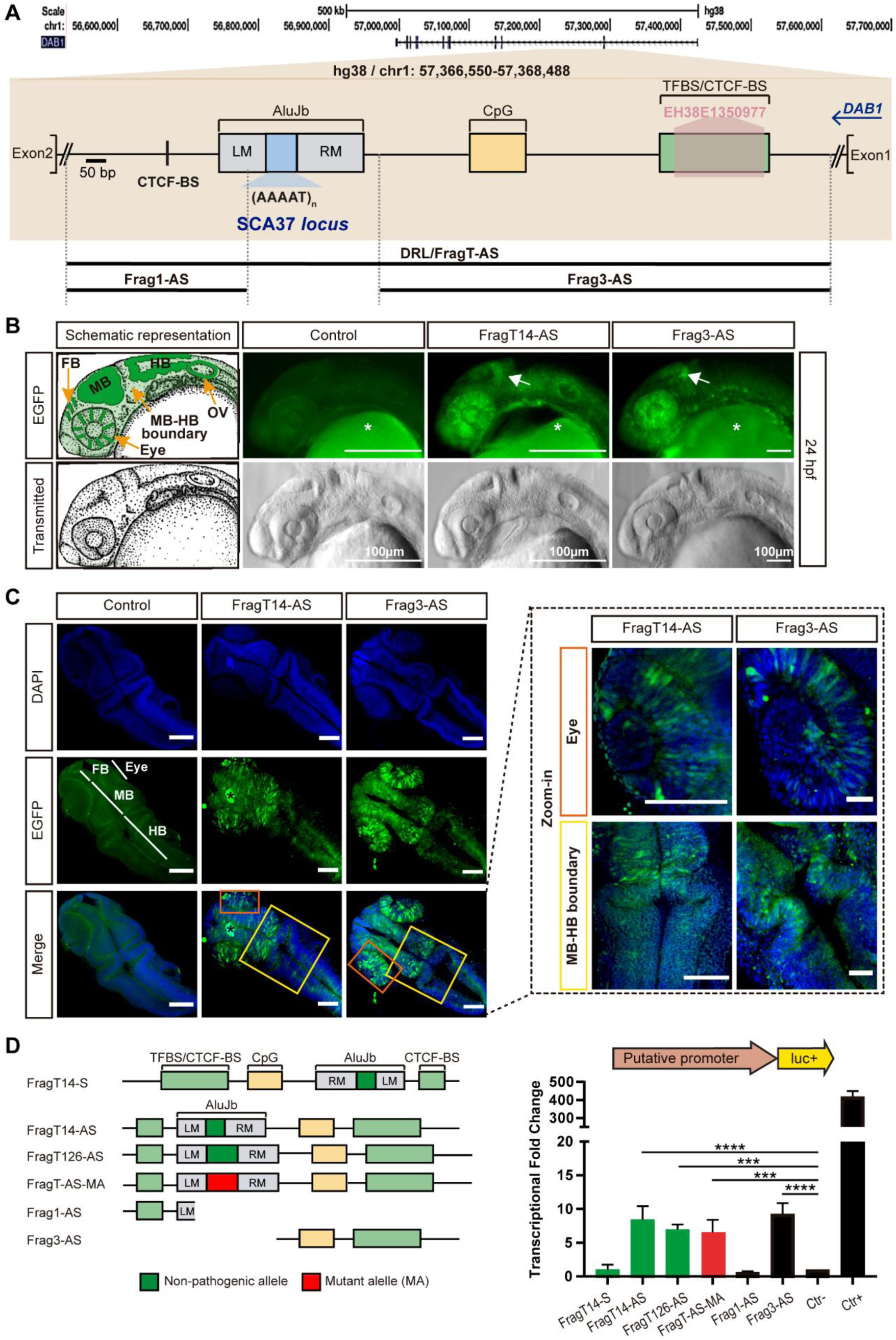
The DRL shows neurodevelopmental enhancer activity. (A) Schematic representation of the DRL/FragT sequence in *DAB1*-antisense strand. The zoom-in box (pale brown box) shows the (AAAAT)_n_ (blue box) in the middle A-rich region of the AluJb element with left (LM) and right (RM) monomers, close to the CpG island (yellow box) and the TFBS (green box), flanked by CTCF-binding sites. The ENCODE-annotated regulatory EH38E1350977 element is represented in pink. Lines below represent the sequences cloned in the zebrafish transgenic lines generated, across and within the DRL sequence. (B) Stereomicroscopic images of EGFP expression in the eye and whole brain, at 24 hpf, in transgenic lines with FragT14-AS and Frag3-AS sequences cloned upstream of a promoter-less EGFP; no expression is seen in control embryos; images were processed with the Fiji software. (C) Left: representative confocal images confirming FragT14-AS and Frag3-AS expression in eye, forebrain, midbrain and hindbrain, in these zebrafish transgenic lines, at 24 hpf; maximum intensity z-projections of 14 planes; scale bars = 100 µm. Right: zoom-in images for the same zebrafish lines; orange and yellow delimitations represent eye and midbrain-hindbrain boundary close-up, respectively; scale bars = 100 µm. (D) Left: schematic representation of the DRL sequences cloned upstream of promoter-less *luciferase* for reporter assays in human HEK293T cells. Right: graphic representation of transcriptional fold change for each sequence, as luc+/Nluc expression ratios in HEK293T cells, compared to the negative control (data from three biological replicates, as mean ± SD; one-way ANOVA test with Dunnett’s post-hoc test; ****p<0.0001, ***p<0.001). FB, forebrain; MB, midbrain; HD, hindbrain; OV, optic vesicle; scale bars = 100 µm. Asterisks represent autofluorescence regions. See also Figure S4.

### *In silico DAB1* mRNA expression analysis

The expression levels of *DAB1* mRNA variants were obtained from Genotype-Tissue Expression (GTEx) V6 for the different brain regions.

### Hi-C data visualization

To detect the predicted *DAB1* TAD position and architecture, the Hi-C data from human cerebellum were obtained from ENCODE^42^ (Encode DCC 2017). TAD position was visualized, at 10 kb resolution, with the 3D Genome browser^43^, from the genomic positions (hg38) Chr1:56,500,000-59,000,000, followed by TAD borders prediction with the 3D Genome Browser.

### Cells and tissues with histone marks for active chromatin

Cells and tissues with ChIP-seq signal for active chromatin histone marks (H3K27ac; H3K4me3 and H3K4me1) in the hg38 Chr1:57,366,550-57,368,488 genomic region were obtained from the ChIP-Atlas Enrichment analysis tool^44,45^ and visualized on Integrative Genomics Viewer V2.16.

### Cloning of DRL/FragT sequences for *in Vivo* Zebrafish Assays

For cloning of the long ∼2kb Total fragment (FragT14) with an unaffected allele (ATTTT/AAAAT)_14,_ spanning the *DAB1* repeat *locus* region (DRL), the genomic DNA was PCR amplified with 200 ng DNA, 200 µM dNTPs, 1.4 mM MgCl_2_, 60 mM Tris-SO4 (pH 9.1), 18 mM (NH4)2SO4, 2 mM MgSO4, 0.3 µM each Alu-CTCF-F and Alu-CTCF-R primer (Table S2), and 1 µL of Elongase enzyme (#10480-010, Invitrogen, Thermo Fisher Scientific). The DNA was initially denatured at 94 °C for 3 min, followed by 10 cycles of amplification (94 °C for 30 s and 62°C for 8 min) and additional 30 cycles of elongation (94 °C for 30 s and 62 °C for 8 min with increments of 20 s per cycle). The PCR product was separated by electrophoresis in 1% agarose (#GA110, Grisp) gels; each DNA band was extracted from the gel with QIAquick Gel Extraction Kit (#28704, QIAGEN), followed by A-overhangs treatment with 1U Taq DNA polymerase (#10342020, Invitrogen, Thermo Fisher Scientific) 1x Taq polymerase buffer with Mg^2+^, 200 µM dNTPs, in 50 µL reaction, at 72 °C for 20 min. This FragT14 sequence, containing the non-pathological allele, was cloned in the entry vector pCR8/GW/TOPO TA (#K250020, Invitrogen, Thermo Fisher Scientific) following the manufacturer’s instructions. FragT with pathogenic sequence (FragT-MA) was amplified as described (in “DRL/FragT activity in HEK293T cells by dual luciferase assays” section). Frag1, Frag2 and Frag3, were amplified from pCR8/GW/TOPO TA (#K250020, Invitrogen, Thermo Fisher Scientific) containing FragT14, using 0.3-0.4 ng of DNA, 1x PCR buffer with Mg^2+^, 1 mM dNTPs, 2.5 U of i-Max II DNA polymerase (#25261, iNtRON Biotechnology), 0.3 µM of primers Alu-Frag1-F and Alu-Frag1-R1 for Frag1 (Table S2); Alu24F and Alu24R for Frag 2 (Table S2) and Alu-Frag2-F and Alu-Frag2-R for Frag3 (Table S2), in 20 µL. The DNA underwent an initial denaturation at 94 °C for 3 min, followed by 30 cycles of 30 s at 94 °C; 4 s at 56 °C and 50 s at 72 °C, and a final extension of 7 min at 72 °C, followed by cloning in pCR8/GW/TOPO TA (#K250020, Invitrogen, Thermo Fisher Scientific) entry vector. These sequences (FragT14, Frag1, Frag2 and Frag3) were cloned in this entry vector in both *DAB1* sense and antisense orientations. These entry vectors were used to recombine each insert into the required destination vector using Gateway LR Clonase II Enzyme mix (#11791020, Invitrogen, Thermo Fisher Scientific) following the manufacturer’s instructions. Each insert sequence was confirmed by Sanger sequencing in the destination plasmid vector. Prior to zebrafish microinjection, each plasmid vector was purified with DNA Clean & Concentrator kit (#D4013, ZYMO Research) according to the manufacturer’s recommendations and resuspended in nuclease-free water.

### *In vivo* DRL/FragT activity in zebrafish

For *in vivo* DRL activity, pCR8/GW/TOPO TA (#K250020, Invitrogen, Thermo Fisher Scientific) entry vector containing FragT14, Frag1, Frag2 and Frag3 sequences (see “Cloning of DRL/FragT sequences for *in vivo* zebrafish assays”) were recombined with the destination *Tol2* promoter-less test vector with an EGFP reporter^46^. Transgenic zebrafish stable lines were established for FragT14-AS, Frag1-AS and Frga3-AS sequences. DRL activity in the brain and eye was assessed at 24 hours postfertilization (hpf) by means of EGFP fluorescence. Embryos were fixed with 4% PFA (#GF750170, Delta Microscopies) in PBS (137 mM NaCl (#31434-M, Sigma-Aldrich, Merck), 2.7 mM KCl (#P9541, Sigma-Aldrich, Merck), 10 mM Na2HPO4 (#A2943, PanReac AppliChem ITW Reagents), 1.8 mM KH2PO4 (#529568, Sigma-Aldrich, Merck)) for 3 h at 4°C, permeabilized with 0.5% Triton X-100 (#T8787, Sigma-Aldrich, Merck) in PBS for 30 min, at room temperature and incubated with DAPI (#D9542, Sigma-Aldrich, Merck). Images were acquired in an inverted Leica SP8 single point scanning confocal microscope with HC PL APO 20x/0.75 IMM/CORR CS2 objective (Leica Microsystems). Eye and brain expression of the DRL sequences was assessed at 24 hpf, while muscle expression was assessed at 72 hpf using Leica M205 FA stereomicroscope using Orca Flash 4.0 LT system (Hamamatsu Photonics). For better visualization, stereomicroscope images of eye and brain expression were processed on Fiji^47^ software, as follows: subtraction of background (50 pixels), deletion of yolk region, subtraction of background (50 pixels), 1.25x multiplication of each pixel and Gaussian filter (2.0 pixels).

### *In vivo* DRL/FragT Interaction with Z48 midbrain enhancer

To study the interaction ability of the DRL region with the midbrain Z48 enhancer, pCR8/GW/TOPO TA (#K250020, Invitrogen, Thermo Fisher Scientific) entry vectors containing the sequences, in *DAB1* antisense orientation, FragT14-AS and the fragmented Frag1-AS and Frag3-AS (see “Cloning of DRL/FragT sequences for *in vivo* zebrafish assays” section) were *in vitro* recombined with a destination vector containing the entry site followed by EGFP^46^ and the midbrain-specific enhancer, Z48^48^. As negative control, the same vector was used without any sequence. Each construct was co-injected with the *Tol2* transposase mRNA in zebrafish embryos, in three independent sessions, with at least 200 embryos per condition. EGFP expression in zebrafish muscle fibers and midbrain was analyzed in all F0 mosaic embryos, at 48 hpf, using a Leica M205 FA stereomicroscope (Leica Microsystems) equipped with an Orca Flash 4.0 LT system (Hamamatsu Photonics).

### *In vivo* silencer assay

The *in vivo* silencer activity of the DRL region was assessed using the *Tol2* vector designated pminiTol2-Z48-CARGFP vector^49^. The sequences to be tested for silencer activity were recombined from the entry vectors (see “Cloning of DRL/FragT sequences for *in vivo* zebrafish assays”) to this vector, between a midbrain-specific enhancer (Z48) and the cardiac actin promoter driving EGFP expression in muscle (transgenesis control). In the empty vector, interaction between Z48 and the cardiac actin promoter drives EGFP to somites and midbrain. Each of these constructs was co-injected with the *Tol2* transposase mRNA in one-to two-cell stage zebrafish embryos, performing three independent microinjection sessions, with at least 100 embryos per condition. Then, 20 embryos were imaged, per condition, at 24 hpf, using a Leica M205 FA stereomicroscope (Leica Microsystems). The EGFP expression was analyzed with Fiji^47^ software, using a custom-built macro to read lif files and save each series as tiff file and another to quantify EGFP intensity in zebrafish somites and midbrain, code available at https://doi.org/10.5281/zenodo.14963259.

### *In vivo* characterization of DRL/FragT enhancer activity

For transient *in vivo* enhancer assays, the above generated pCR8/GW/TOPO TA (#K250020, Invitrogen, Thermo Fisher Scientific) entry vector containing, in *DAB1* orientation, the non-pathogenic (FragT14-S), and the pME-MCS with pathogenic FragT (FragT-S-MA) were recombined with the ZED transgenesis vector^49^. Mosaic transgenic zebrafish were generated by Tol2-mediated transgenesis^37^. Embryos showing expression of mCherry in the muscle fibers (transgenesis control) were selected for enhancer assessment, at 4 days postfertilization (dpf). Then, larvae were analyzed for the presence of EGFP-positive cells in the cerebellum or in the forebrain, using an inverted Leica SP5 single point scanning confocal microscope (Leica Microsystems) with a HC PL APO Lbl. Blue 20x/0.70 IMM/CORR objective. These images were analyzed and quantified using Fiji^47^ and Imaris (version 10.2.0, Andor Oxford) software. One larva was considered positive if at least one EGFP-positive cell was detected within the cerebellum or forebrain domains. Data were presented as percentage of positive larvae to ensure the quantification of transposon integration in different embryos.

### Cell Transfection

HEK293T cells were seeded in 6-well cell culture plates at a density of 120,000 cells per well and transfected with jetPRIME transfection reagent (#1011000027, Polyplus). hNSC were plated in 24-well cell culture plates at a density of 70,000 cells and transfected with Lipofectamine 2000 (#11668030, Thermo Fisher Scientific).

### Cloning of sequences in DRL/FragT region for cell assays

The DRL/FragT, in *DAB1* sense and antisense orientations, containing the non-pathogenic repeat with 126 units (FragT126) or SCA37 pathogenic (FragT-MA) allele, was amplified from genomic DNA. They were amplified by Long-Range PCR, using 1 ng of each vector, 200 µM each dNTP, 1x PrimeSTAR GXL Buffer Mg2+ Plus, 1.25 U PrimeSTAR GXL DNA polymerase (#R050A; Takara Bio), 10 µM primers containing EcoRI or XmiI tails (sense: Alu-CTCF-2F-EcoRI, Alu-CTCF-2R-XmiI; antisense orientation: Alu-CTCF-2F-XmiI and Alu-CTCF-2R-EcoRI; Table S2) and water up to 50 µL. After 1 minute of initial denaturation at 98 °C, DNA samples underwent 30 cycles of amplification (98 °C for 10 min and 68 °C for 5 min). PCR products were separated by agarose gel electrophoreses and the DNA band of interest purified with Zymoclean Gel DNA Recovery kit (#D4008; ZYMO Research) and digested overnight with EcoRI (#IVGN0116, Anza, Thermo Fisher Scientific) and XmiI (#IVGN0966, Anza, Thermo Fisher Scientific), at 37 °C. Restriction products were purified using the DNA Clean & Concentrator kit (#D4013, ZYMO Research) and cloned into the Tol2kit entry vector pME-MCS^50^, previously linearized with EcoRI (#IVGN0116, Anza, Thermo Fisher Scientific) and XmiI (#IVGN0966, Anza, Thermo Fisher Scientific), at 37 °C. The non-pathogenic (FragT14) was cloned as described (see “Cloning of DRL/FragT sequences for *in vivo* zebrafish assays”). FragT14-S sub-fragments Frag1-S, Frag2-S, Frag3-S, Frag1+Frag2-S, Frag2+CpG-S and Frag2+Frag3-S were amplified from the generated pCR8/GW/TOPO TA (#K250020, Invitrogen, Thermo Fisher Scientific) entry vector containing FragT14-S, using primers detailed (Table S2). Frag2-S-MA was amplified from pME-MCS containing FragT-S-MA, by Long-Range PCR using primers containing 5’-tails compatible with EcoRI and XmiI restriction enzymes (Alu24F_XmiI and Alu24R_EcoRI; Table S2). These subfragments, Frag1-S, Frag2-S, Frag3-S, Frag1+Frag2-S, Frag2+CpG-S and Frag2+Frag3-S were inserted into the pCR8/GW/TOPO TA (#K250020, Invitrogen, Thermo Fisher Scientific), whereas Frag2-S-MA was cloned into pME-MCS vector. They were recombined with the required destination vector using Gateway LR Clonase II Enzyme mix (#11791020, Invitrogen, Thermo Fisher Scientific) and each recombined sequence was confirmed by Sanger sequencing.

### FragT/DRL activity in HEK293T cells by dual luciferase assays

To generate the pGL3/GW vector, the Gateway (GW) sequence was digested from pGL4.23/GW vector (#60323; Addgene)^51^, and inserted in pGL3-basic luciferase reporter vector (#E175A, Promega) using KpnI (#IVGN0178, Anza, Thermo Fisher Scientific) and BglII (#IVGN0198, Anza, Thermo Fisher Scientific) restriction sites. For activity test in cells, the pCR8/GW/TOPO TA (#K250020, Invitrogen, Thermo Fisher Scientific) vectors (from “Cloning of sequences in DRL/FragT region for cell assays” section) containing the FragT14-S and FragT14-AS, Frag1-AS, and Frag3-AS were recombined with the pGL3-basic/GW. These pGL3-basic luciferase reporter test vectors were co-transfected with pNL1.1TK[Nluc/TK (#N1501; Promega), in HEK293T cells. The activity of these sequences was assessed 24 hours post-transfection, by Dual Luciferase Assay using Nano-Glo® Dual-Luciferase® Reporter Assay System (#N1610, Promega). In each well, the firefly *luciferase* (Fluc) (encoded by *luc2*) activity was measured, followed by the NanoLuc® (Nluc) activity to normalize luminescence levels. Results were presented as luc2/Nluc ratios, relative to pGL3-basic empty (negative control). Three independent transfection replicates and three technical replicates were carried out for each luciferase activity measurement. Luminescence produced was measured in the Synergy 2 plate reader (BioTek), using the Gen5™ software.

### *In vitro* silencer assays

The thymidine kinase (TK) promoter was cloned in pGL4.23 GW[luc2/minP] (#60323; Addgene)^51^ downstream the GW sequence using XhoI (#IVGN0086, Anza, Thermo Fisher Scientific) and BglII (#IVGN0198, Anza, Thermo Fisher Scientific) restriction enzymes to generate the luciferase reporter vector pGL4.23 GW[luc2/Tk] vector. The fragments in entry vectors (from “Cloning of sequences in DRL/FragT region for cell assays” section) were recombined into the pGL4.23 GW[luc2/Tk]. The generated constructs were co-transfected along with pNL1.1TK[Nluc/TK] (#N1501, Promega) in HEK297T cells, plated in a 24-well plate, using jetPRIME transfection reagent (#101000027, Polyplus). Luciferase activity measurements, assessed in triplicate in three independent transfection replicates, and statistical analysis were performed as described in section “FragT/DRL activity in HEK293T cells by dual luciferase assays”.

### Enhancer assays in human neural stem cells

The fragments in entry vectors (from “Cloning of sequences in DRL/FragT region for cell assays” section) were recombined into the destination luciferase reporter vector pGL4.23 GW[luc2/minP] (#60323; Addgene)^51^. The generated enhancer luciferase reporter constructs were co-transfected along with pNL1.1TK[Nluc/TK (#N1501; Promega) in hNSC plated in 24-well format, using Lipofectamine 2000 (#11668030, Thermo Fisher Scientific), following manufacturer’s instructions. The positive control (pGL4.23 [luc2/Tk])^52^ containing the tyrosine kinase promoter was generated from pGL4.23 GW[luc2/minP]. To use as a negative control, a genomic region without histone marks was cloned into the pGL4.23 GW [luc2/minP] vector ^53^. Luciferase activity assessment was performed in three technical replicates as described above, in three independent replicates of the transfection.

### Characterization of *DAB1* mRNA variants expression in hNSC

The expression of total *DAB1* mRNA and *DAB1* mRNA variants in hNSC was confirmed by semiquantitative PCR using the primers described (Table S2). cDNA was synthesized from 1 µg total RNA of hNSC or human cerebellum (#636535, Clontech), using SuperScript III First-Strand Synthesis SuperMix for qRT-PCR (#11752050, Invitrogen, Thermo Fisher Scientific), following the manufacturer’s procedure. cDNA was amplified by standard PCR using 6.25 µM of each primer, 1x Supreme NZYTaq II Green Master Mix (#MB36001, NZYtech), and water up to 50 µL. For PCR reaction, an initial denaturation, at 95°C for 5 min, was followed by 32 cycles of amplification (94 °C 30 s, 60 °C 30 s and 72°C 30 s) and a final extension, at 70°C for 10 min.

### Immunofluorescence

For iPSC lines, pluripotency markers OCT4, SOX2, TRA-1-60 and SSEA4 expression was confirmed with rabbit OCT4, rat SOX2, mouse SSEA4 and mouse TRA-1-60 primary antibodies combined with Alexa Fluor 594 Donkey anti-Rabbit, Alexa Fluor 488 Donkey anti-Rat, Alexa Fluor 488 goat anti-mouse and Alexa Fluor 594 goat anti-mouse secondary antibodies, using PSC 4-Marker Immunocytochemistry Kit (#A24881, Thermo Fisher Scientific), following the manufacturer’s instructions. After immunocytochemistry, iPSCs were imaged on Axio Observer.Z1 (Zeiss) with a 10x objective. The expression of protein DAB1 and the neuronal markers TUJ1 and MAP2 was confirmed in iPSNs by immunofluorescence. iPSNs were fixed in 4% PFA (#GF750170, Delta Microscopies) in PBS, for 10 min, at room temperature, permeabilized in 0.2% Triton X-100 (#T8787, Sigma-Aldrich, Merck) for 10 min, at room temperature and blocked in 5% BSA (#MB04601, NZYtech) for 30 min, at room temperature. Cells were incubated with mouse β-TuJ-1 (1:500, #MAB1195, R&D Systems) and mouse MAP2 (1:250, #MAB3418, Sigma-Aldrich, Merck) primary antibodies diluted in blocking solution, overnight, at 4°C. iPSN cells were then incubated with secondary goat anti-mouse antibodies Alexa Fluor 488 (1:1000, #A-10680, Thermo Fisher Scientific) and Alexa Fluor 594 (1:1000, #A-11005, Thermo Fisher Scientific), for 1 h, at room temperature. Cells were mounted in ProLong Gold with DAPI (#P36931, Invitrogen, Thermo Fisher Scientific) and imaged on Axio Observer.Z1 (Zeiss) with Planapo 63x/1.4 oil objective. DAB1 protein expression in hNSC was also confirmed by immunofluorescence with rabbit polyclonal anti-DAB1 antibody. Cells were fixed with 4% PFA (#GF750170, Delta Microscopies) in PBS, 48 h post-transfection. After permeabilization with 0.2% Triton X-100 (#T8787, Sigma-Aldrich, Merck) and blocking with 5% BSA (#MB04601, NZYtech), cells were incubated overnight, at 4 °C, with rabbit polyclonal anti-DAB1 antibody (1:100, sc-13981, Santa Cruz Biotechnology). On the following day, cells were incubated with Alexa Fluor 594 goat anti-rabbit (1:1000, #A-11012, Thermo Fisher Scientific) for 1 h at room temperature, counterstained with DAPI (#D9542, Sigma-Aldrich, Merck) and analyzed on a Axio Observer.Z1 (Zeiss) with Planapo 63x/1.4 oil objective.

Whole-mount immunohistochemistry of zebrafish embryos and larvae was performed at 24 hpf and 5 dpf. For primary motor neuron (PMN) axon staining, 24 hpf WTAB zebrafish embryos, previously injected with 100 ng/µL *DAB1* mRNA, were fixed for 3 h at 4°C in 4% PFA (#GF750170, Delta Microscopies) in PBS with 0.1% Triton X-100 (#T8787, Sigma-Aldrich, Merck) (PBS-T). Permeabilization was performed with 1% Triton X-100 (#T8787, Sigma-Aldrich, Merck) in PBS, for 2 h at room temperature and the blocking was carried out using 1% BSA (#MB04601, NZYtech), 1% DMSO (#D8418, Sigma-Aldrich, Merck), 1% NGS (#X0907, Dako), 0.7% Triton X-100 (#T8787, Sigma-Aldrich, Merck) in PBS for 1 h at room temperature. Embryos were incubated with anti-synaptic vesicle 2 (SV2) antibody (1:200, #AB2315387, Developmental Studies Hybridoma Bank) diluted in blocking solution overnight, at 4°C. Embryos were washed and incubated with Alexa Fluor-647 goat anti-mouse IgG antibody (1:750, #A-21235, Thermo Fischer Scientific) and DAPI (1:1000; #D9542, Sigma-Aldrich, Merck) diluted in blocking solution for 4 h, at room temperature. After washing solutions, embryos were mounted in microscope slides using glycerol-based ibidi mounting media (#50001, ibidi). The axonal length of GFP+ cerebellar granule cells was analyzed in larvae from transgenic line gSA2AzGFF152B; UAS:GFP^33^ (kindly shared by Drs. Kawakami and Hibi, Japan), at 5 dpf, microinjected with 200 ng/µL *DAB1* mRNA. Larvae were fixed as above, permeabilized with 0.5% Triton X-100 (#T8787, Sigma-Aldrich, Merck) in PBS, for 30 min, at room temperature, and incubated with DAPI (1:1000, #D9542, Sigma-Aldrich, Merck) overnight, at 4°C. In the next day, larvae were washed with PBS-T and mounted on microscope slides using glycerol-based ibidi mounting media (#50001, ibidi). Axon length analysis was carried out as described in “Analysis of axonal tracts in zebrafish”.

### CRISPR/Cas9 deletion of FragT/DLR

#### Synthesis of sgRNA encoding plasmids

Small guide RNAs (sgRNAs) predicted to target and delete FragT/DRL by CRISPR/Cas9, suitable to use with S. Pyogenes Cas9 enzyme, were selected from the CRISPR targets Track of the UCSC genome browser. Two sgRNAs targeting the region flanking the *DAB1* (ATTTT)_n_ were chosen, one upstream and one downstream of the FragT/DRL, with a MIT specificity score higher than 50 and a Doench efficiency score higher than 55^54^. Oligonucleotides encoding the selected sgRNAs with overhangs compatible with BpiI restriction sites (Table S2) were annealed *in vitro* in Annealing buffer (5mM Tris (#MB01601, NZYtech) pH7.5-8, 25mM NaCl (#MB15901, NZYtech) and 0.5mM EDTA (#20302, VWR) by incubation at 95°C for 5 min, followed by a cool down until 85°C at 2°C/s and from 85°C to 25°C at 0.1 °C/s. Annealed oligos targeting the FragT upstream region (sgCTCF1_U1; Table S3) were cloned in pSpCas9(BB)-2A-GFP (PX458) vector (#48138, Addgene) and the annealed oligos targeting the FragT downstream region (sgCTCF2_1.1; Table S3) were inserted in pU6-(BbsI) CBh-Cas9-T2A-mCherry (#64324, Addgene) plasmid using T4 DNA ligase (#10481220001, Sigma-Aldrich, Merck), after linearization with BpiI (#IVGN0044, Anza, Thermo Fisher Scientific) restriction enzyme.

#### CRISPR/Cas9 deletion

HEK293T and hNSC cells were co-transfected with plasmid vectors encoding the sgRNAs. Co-transfected HEK293T cells were isolated by fluorescence-activated cell sorting (FACS), 48 h post-transfection, dropping a single cell per well in a 96-well plate. HEK293T single cells were cultured until colony formation. For each hNSC transfection event, the co-transfected and non-transfected cells from the same culture plate were isolated by FACS, in a FACSARIA II cell sorter (BD Biosciences), 48 h post-transfection. DNA was extracted from HEK293T after clone formation and hNSC sorted cells using the Quick-DNA Miniprep kit (#D4068S, ZYMO Research). Cells were genotyped by conventional PCR using primers Alu-CTCF-2F and Alu-CTCF-2R (Table S2) to confirm DNA deletion of FragT/DRL (ΔFragT), of ∼2 Kb. HEK293T clones and hNSC sorted cells from co-transfection of plasmid vectors encoding sgCTCF1_U1 and sgCTCF2_1.1 produced homozygous deletions and were selected for RNA extraction.

### RNA Extraction from CRISPR/Cas9 deleted cells

Total RNA in HEK293T cells was isolated from clones homozygous for the deletion, ΔFragT, using the RNeasy minikit (#74104, Qiagen) following the manufacturer’s procedure. Total RNA from sorted double-stained (GFP^+^ and mCherry^+^) and unstained hNSC was isolated with PureLink™ RNA Micro Scale Kit (#12183016, Invitrogen, Thermo Fisher Scientific) according to manufacturer’s instructions.

### Analysis of Gene Expression by Real-time Quantitative RT-PCR

The total RNA isolated from ΔFragT (Crispant) hNSC and HEK293T and their respective non-deleted controls was retrotranscribed into cDNA using SuperScript III First-Strand Synthesis SuperMix for qRT-PCR (#11752050, Thermo Fisher Scientific), following the manufacturer’s procedure. The real-time quantitative RT-PCR (qRT-PCR) reaction for each mix to amplify *DAB1*, *DAB1* variants (V1 and V3; V2; and V4), *MYSM1*, *OMA1* and *FYB2* was prepared in a total volume of 10 µL, containing 1x iTaq SYBR Green Supermix, 0.25 or 0.50 µM of each forward and reverse primers for HEK293T or hNSC cDNAs, respectively, 1 µL of cDNA (diluted 1:10), and nuclease-free water up to the final volume. The reactions were performed on a CFX96 Touch Real-Time PCR Detection System (Bio-Rad). The program included an initial denaturation step of 95°C for 3 minutes, followed by 40 cycles of 10-second denaturation at 95°C and 30-second annealing at 60°C, with fluorescence acquisition at the end of each cycle. The melt curve analysis was conducted by doing a step of 65°C to 95°C with 0.5°C increments of 10 seconds, to assess the specificity of the amplified products. Total *DAB1*, *DAB1* variants (V1 and V3; V2; and V4), *MYSM1*, *OMA1* and *FYB2* mRNA levels were then quantified using 2^ΔΔct^ and expression normalized with *GNA13* and *CAPN10* reference genes (Table S2). For hNSC and HEK293T CRISPR cells, three biological replicates were used. Three technical replicates were carried out in qRT-PCR for each sample.

### Cap analysis of gene expression (CAGE)

Cap analysis of gene expression (CAGE) was performed in iPSN derived from one or two clones of three SCA37 iPSC affected (AF2Cl4; AF2Cl21; AF3Cl7 and AF4Cl32) and three unaffected individuals (GM23280; ND41866 and NAS6). Prior to CAGE library generation, total RNA was isolated from iPSN at day 8 of differentiation with TRIzol reagent (#15596026, Thermo Fisher Scientific), followed by purification with RNeasy mini KIT (#74104, Qiagen), according to the manufacturer’s protocol. The integrity and concentration of the RNA were assessed using the Agilent TapeStation system. CAGE libraries were constructed as previously described^55^. Briefly, complementary DNA (cDNA) was synthesized from total RNA using random primers and reverse transcriptase. The 5’ end of mRNA was specifically captured using the cap-trapper method and ligated to a linker. The captured RNA was then subjected to first-strand cDNA synthesis, and second-strand synthesis was performed to generate double-stranded cDNA. This cDNA was then digested with EcoP15I (#R0646S, NEB) to generate short tags, that were ligated to adapters and amplified by PCR, purified and sequenced on the Illumina HiSeq 2500 generating 50 bp single-end reads at an average of 50M reads/sample. Illumina reads were demultiplexed and trimmed using the FASTX toolkit (hannonlab.cshl.edu/fastx_toolkit/), to remove adapter sequences and low-quality bases. CAGEseq reads were filtered for artifacts using TagDust (version 1.12) and mapped to the human genome (hg19), using BWA for short reads (version 0.5.9). Mapped CAGEseq reads were processed and organized into CAGE-clusters using a series of Python scripts developed at the RIKEN Omics Science Center^55^. In summary, single-base pair promoters were identified by locating all genomic positions corresponding to the 5’ end of at least one CAGEseq read, excluding those with a mapping quality below 20 to filter out multimapping reads. Single base pair promoters within 20 bp of each other were grouped into a single CAGE-cluster. Raw counts were then normalized by dividing the number of CAGEseq reads at each CAGE-cluster by the total number of mapped tags in the library, and multiplying by one million (tags per million, TPM). CAGE clusters were annotated using GENCODE v17, named using the gene symbol they mapped to and visualized on the ZENBU omics data integration and visualization system^56^. Gene ontology analysis was performed using the functional annotation tool of Database for Annotation, Visualization and Integrated Discovery (DAVID)^57^ (https://david.ncifcrf.gov/), for a total of 447 genes found differentially expressed between SCA37 affected and unaffected individuals, with p-value <0.05 and - 1.5<Log_2_(FC)>1.5.

### Analysis of axonal tracts in zebrafish

To assess primary motor neurons (PMN) axons, at 24 hpf, zebrafish embryos were injected with *DAB1* mRNA, and immuno-stained with SV2 antibody (see “Immunofluorescence” section). The 6-somites region spanning the cloaca of four replicates with at least five zebrafish injected embryos per condition was imaged on an inverted Leica SP8 single point scanning confocal microscope (Leica Microsystems) with a PL APO 40x/1.10 water CS2 objective. PMN axon length was measured in maximum intensity z-projections with Fiji^47^ NeuroJ^58^ plugin.

To analyze the axonal length of GFP+ cerebellar granule cells at 5 dpf, zebrafish larvae from gSA2AzGFF152B; UAS:GFP transgenic line^33^ were microinjected with *DAB1* mRNA, and whole-mounted staining was performed as described in the “Immunofluorescence” section. Three replicates with at least 20 larvae/condition were imaged on an inverted Leica SP8 single point scanning confocal microscope with a HC PL APO 10x/0.40 CS2 dry objective with a zoom of 2.2 (Leica Microsystems), and three-dimensional imaging analysis was performed in Imaris (version 10.2.0, Andor Oxford) software.

### Protein extraction and western blot

For human DAB1 protein expression in zebrafish, 217 embryos of each condition (control and DAB1 mRNA injection) were deyolked at 24 hpf with Ginzburg Fish Ringers solution (55 mM NaCl (#S/3161/60, Fisher Chemical), 1.8 mM KCl (#2676.298, VWR), 1.25 mM NaHCO3 (#S5761, Sigma-Aldrich, Merck)) and homogenized on ice in radioimmunoprecipitation assay (RIPA; #R0278, Sigma-Aldrich, Merck) buffer supplemented with protease inhibitor cocktail (#11873580001, Roche, Merck) and phosphatase inhibitor (#S6508, Sigma-Aldrich, Merck). Samples were incubated on ice for 30 min, centrifuged and sonicated. Protein quantification was assessed using Pierce BCA Protein Assay kit (#23227, Thermo Fisher Scientific). Protein lysates (20 µg per condition) were denatured at 95°C for 5 min, resolved on a 10% SDS-PAGE gel and transferred to 0.2 µm nitrocellulose membrane (#10600001, GE Healthcare) overnight at 4°C, using a wet transfer system. Membrane was washed in PBS with 0.1% Tween-20 (PBST, #P9416, Sigma-Aldrich, Merck), blocked with 5% nonfat dry milk in PBST for 1 h and incubated with rabbit anti-DAB1 antibody (1:500, sc-13981, Santa Cruz Biotechnology) overnight at 4°C. Membrane was washed and incubated with anti-rabbit horseradish peroxidase (HRP)-conjugated antibody (1:10000, #111-035-003, Jackson ImmunoResearch) for 2 h. Then, it was washed, and Clarity Western ECL Substrate (#1705060, Bio-Rad) was added to the membrane for protein visualization by chemiluminescence on a ChemiDoc XRS+ (Bio-Rad). Membrane was stripped, re-probed with mouse anti-GAPDH antibody (1:20000, #HRP-60004, ProteinTech Group) overnight at 4°C and incubated with anti-mouse HRP-conjugated antibody (1:10000, #711-035-151, Jackson ImmunoResearch), as above for protein loading control. As positive control, total protein extracts were extracted from HEK293T cells transfected with pCMV6-XL4-DAB1 mammalian expression vector (#SC113027, Origene)^8^ and 1 µg was loaded on SDS-PAGE gel.

### Statistical analysis

Graphical representation was carried out using GraphPad Prism, and statistical analysis was performed using IBM SPSS. Normal distribution of the data and homogeneity of variances were assessed by Shapiro-Wilk and Levene’s tests, respectively. Details of statistical analyses performed in this manuscript can be found in the figure legends and main text.

## Results

### The SCA37 Repeat Disrupts a *DAB1* Cis-regulatory Element

To investigate how the (ATTTC)_n_ in the DRL region causes SCA37, we started by characterizing the regulatory landscape of *DAB1* in human cerebellar cells. We analyzed genome-wide chromosome conformation capture (Hi-C) data obtained from the ENCODE Project^42^. This showed that *DAB1*, which spans more than 1.2 Mb, is highly structured within a topological domain (TAD) visualized with the 3D Genome Browser^43^ (hg38, Chr1:56,800,000-58,700,000) containing also upstream more centromeric (*OMA1*, *TACSTD2 and MYSM1*) and downstream further telomeric (*C8A* and *C8B*) genes (Figure 1A). Therefore, this TAD likely contains the cis-regulatory elements (CREs) responsible for *DAB1* transcriptional regulation in the human cerebellum. Furthermore, our previous work has shown that the *DAB1* mRNA variants V1, V2, V3 and V4 (Figure 1A) are transcribed through the intronic SCA37 repeat, in the cerebellum^8^. Cap analysis of gene expression (CAGE) from human adult tissue^8^ and transcriptome data from public databases (GTEx) showed that the V1, V3 and V4 are considerably more expressed in cerebellum than in other brain regions (Figure 1B), suggesting that the promoters controlling their expression could be under the influence of CREs that contribute to the dysregulation of *DAB1* in SCA37. To examine this further, we reprogrammed SCA37 fibroblasts into induced pluripotent stem cells (iPSC) (Figure S1). Then, we used these reprogrammed cells to perform circularized chromosome conformation capture sequencing (4C-seq)^38^, setting as viewpoints the promoters of V1, V3 and V4 transcripts and that of the V2 *DAB1* transcript variant, to quantify their spatially most proximal DNA interactions. We found that the promoter of the V2 transcript, that showed low expression in the cerebellum (Figure 1B), interacted mainly with sequences located in the most upstream regions of the *DAB1* regulatory landscape, while the promoters of V1, V3 and V4 interacted mostly with downstream regions, including the DRL sequence containing the *DAB1* ATTTT/AAAAT repeat associated with SCA37 (Figure 1C and Figure S2). To further confirm the interaction of V1, V3 and V4 promoters with the DRL, we performed 4C-seq using the DRL as viewpoint; we detected an interaction with the V1, V3 and V4 promoters, but not with the V2 promoter (Figure 1C and Figure S2). These results indicate that the cerebellar-specific expression of the *DAB1* V1, V3 and V4 variants is defined by specific chromatin contacts, including the DRL sequence. Furthermore, the DRL region in SCA37 individuals, that contains the pathogenic (ATTTC)_n_, seems to have a slight wider range of interactions within the 3’ *DAB1* region, supporting the idea that the DRL genomic sequence has cis-regulatory functions in the control of chromatin architecture and transcription (Figure 1C and Figure S2).

Enhancers are CREs that can activate transcription of distal and nearby promoters^59^. H3K27ac is one of the epigenetic marks associated with active enhancers, while H3K4me1 is usually associated with active and poised enhancers^60^. To find out whether the DRL region has a function in transcriptional regulation during development and in the adult cerebellum, we searched for epigenetic histone modifications associated with active enhancers in cerebellum, iPSC, neural progenitor cells and embryonic stem cells (ESC). We analyzed chromatin immunoprecipitation followed by sequencing (ChIP-seq) signal associated with active enhancers (H3K27ac and H3K4me1) and promoters (H3K4me3), as enhancers can act as autonomous weak promoters^61^. We observed enrichment of these marks in the DRL sequence, in particular, overlapping a CpG island and predicted transcription factor binding sites (TFBS), including predicted binding sites of CTCF (Figure 1D), known to be involved in chromatin looping and organization.

Together with data from other non-neuronal cells and tissues (Figure S3), these results strongly suggest that the DRL genomic sequence has a role in cis-transcriptional regulation.

### The *DAB1* Repeat Locus (DRL) is a neurodevelopmental enhancer

To determine whether the DRL sequence has regulatory functions and their levels of directionality^61^, we sought to investigate its potential in driving transcription *in vivo.* We cloned this DRL sequence, hereby designated as DRL/Total Fragment (FragT; Figure 2A), upstream of a promoter-less EGFP reporter gene integrated in a zebrafish transgenesis vector^46^, and performed *in vivo* reporter assays in zebrafish. FragT, which contains 14 non-pathogenic AAAAT repeats, was integrated in *DAB1*-sense and antisense orientations. After microinjection in zebrafish embryos, we detected EGFP expression driven by *DAB1*-antisense (FragT14-AS), but not by the *DAB1*-oriented (FragT14-S) sequence (Figure S4A). Notably, FragT14-AS stable transgenic lines of the antisense sequence were able to activate EGFP expression in the eyes, forebrain, midbrain and hindbrain of zebrafish embryos, as early as at 24 hpf (Figures 2B and 2C), recapitulating the expression pattern of zebrafish *Dab1* homologues, at the same developmental stage (Figure S4B)^62^. This was accompanied by muscle cell expression that was clearly evident at 72 hpf (Figure S4C).

Next, we assessed whether the regulatory function of the DRL sequence is partitioned into distinct subdomains. To reach this goal, we fragmented Frag-T (with approximately 2kb) into three sub-fragments, in *DAB1*-antisense orientation. Frag1-AS and Frag3-AS were located upstream and downstream of the pentanucleotide repeat tract, and Frag2-AS spanned the AluJb element containing the AAAAT repeat (Figure 2A). After transgenesis and reporter assays using these three fragments, we observed that Frag1-AS and Frag3-AS sequences were able to drive EGFP transcription (Figures 2B and 2C and Figure S4C), while Frag2-AS was not. Furthermore, the expression pattern driven by these different fragments, at 24 hpf, showed that only the Frag3-AS was able to partially recapitulate the pattern seen for FragT-AS sequence (Figures 2B and 2C and Figure S4C), whereas Frag1-AS showed only expression of EGFP in horizontal myoseptum (Figure S4C), a region crucial for transmitting muscular contractility.

To investigate if the DRL region was able to activate transcription in human tissue, we performed reporter assays in human embryonic kidney (HEK293T) cells, similar to that carried out in zebrafish, by cloning the different fragment sequences fused with a promoter-less firefly *luciferase* reporter gene (Figure 2D). As previously, we cloned FragT, containing 14 non-pathogenic AAAAT repeats, in *DAB1*-sense (FragT14-S) and antisense (FragT14-AS) orientations; then, we transfected HEK293T cells and quantified *luciferase* expression. We observed that, as in zebrafish, the sequence in sense orientation was not able to drive reporter gene expression, while in the antisense orientation was (Figure 2D; 8-fold compared to the negative control; p<0.0001). Additionally, we also observed that a higher copy number of the non-pathogenic repeat, up to 126 units (FragT126-AS), did not change the regulatory output of the DRL (Figure 2D; 7-fold compared to the negative control; p<0.001). We further tested Frag1-AS and Frag3-AS, observing that Frag3-AS was able to trigger expression of *luciferase*, while Frag1-AS was not (Figure 2D). These results recapitulate in human cells what was observed *in vivo*, where Frag3-AS had a broader activity in several tissues, including neural and muscle cells, while Frag1-AS activity was only mildly detected in muscle cells.

Overall, these results strongly indicate that the DRL region is a complex CRE that joins multiple regulatory inputs from the combinatorial action of different functional domains present in the DRL/FragT sequence.

### The DRL is an interaction hub of neural regulatory elements

Because the DRL showed to be composed of different functional elements, some of them harboring putative binding sites for CTCF (Figure 2A), known to be required for efficient chromatin looping^63^, we questioned whether the DRL contributes to the establishment of chromatin interactions with other nearby CREs. To test this, we cloned FragT14-AS, Frag1-AS and Frag3-AS from the DRL region upstream of a promoter-less EGFP, and included a strong midbrain enhancer downstream^48^. In this configuration, if the DRL works as a hub for interactions with other CREs, it is expected that the strong midbrain enhancer triggers EGFP expression in this brain region. We observed that FragT14-AS was able to drive broad and strong expression of EGFP in the midbrain (Figure 3A), much broader and stronger than previously observed in zebrafish embryos with the FragT14-AS alone (Figures 2B and 2C); in contrast, in the somites, EGFP expression was similar to that with FragT14-AS (Figure S4C). Additionally, Frag3-AS showed comparable EGFP expression, but weaker in some of the embryos, suggesting that most of the functional interaction domain is located in this fragment, while Frag1-AS showed decreased ability to interact with the midbrain enhancer, with the interactions being restricted to the apical region of the midbrain (Figure 3A).

**Figure 3.**
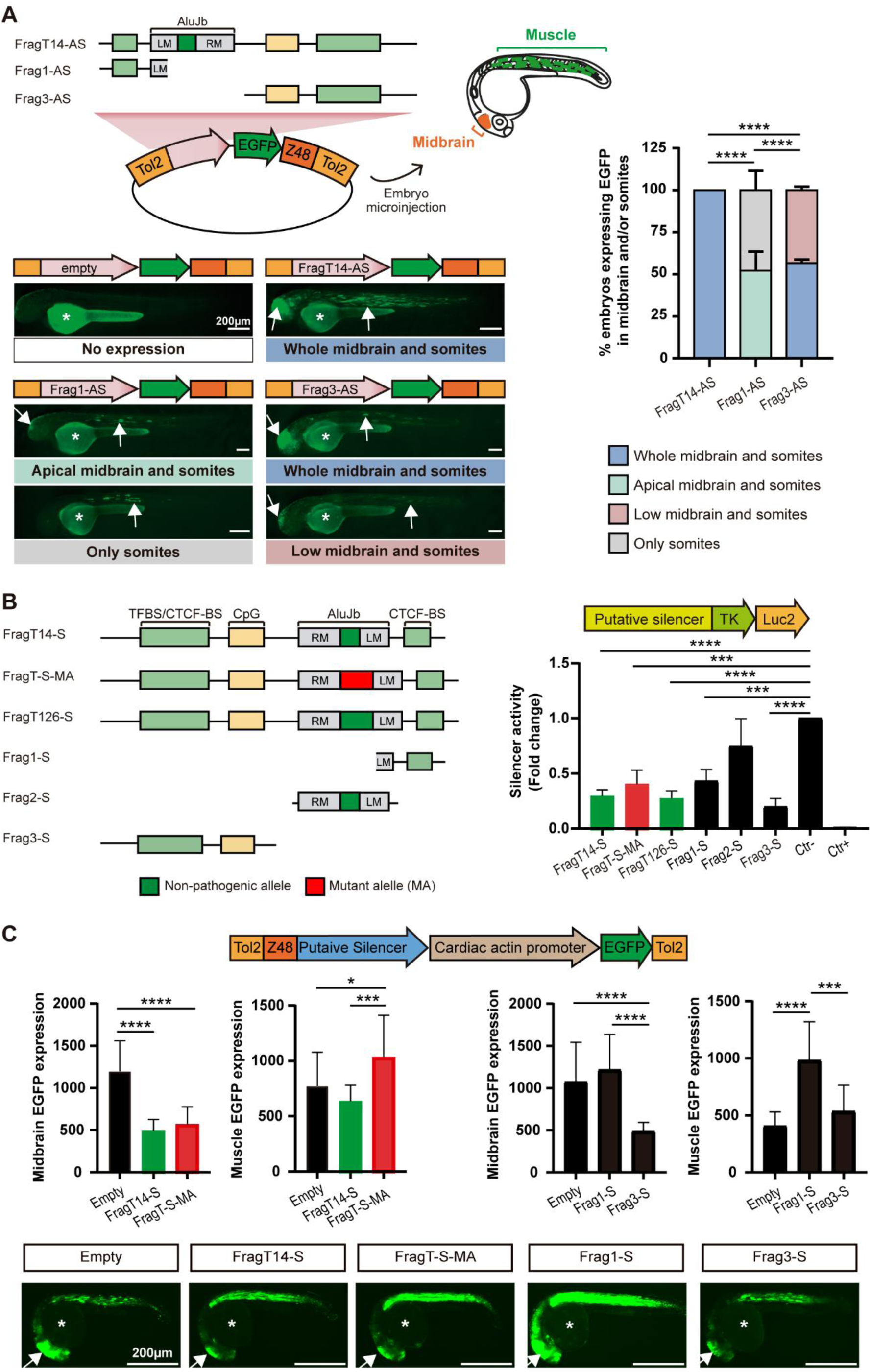
DRL interaction with neural regulatory elements and silencer activity outside of the *DAB1* expression domain. (A) Schematics of interactions between the DRL sequence and the zebrafish midbrain-specific Z48 enhancer, at 48 hpf. Top: FragT14-AS, Frag1-AS and Frag3-AS sequences cloned upstream of a promoter-less EGFP, and Z48 with schematics of their driven EGFP expression to zebrafish somites and upon Z48 interaction only to the whole midbrain. Bottom left: representative stereomicroscopic images of different EGFP expression patterns in midbrain from FragT14-AS, Frag1-AS and Frag3-AS interactions with the Z48 enhancer, in embryos. Bottom right: graphic representation of the percentage of embryos expressing EGFP in midbrain and/or somites (as mean ± SD of three replicates; EGFP positive embryos, FragT14-AS=238, Frag1-AS=195 and Frag3-AS=148, ꭕ^2^, ****p<0.0001). (B) Left: schematic representation of the DRL sequences cloned upstream the TK promoter in the control of *luciferase* expression for silencer assay in HEK293T cells; right: silencer activity in HEK293T cells, measured by the fold chance in luc2/Nluc expression ratios compared to the negative control (mean ± SD of three biological replicates; one-way ANOVA test with Dunnett’s post-hoc test; ****p<0.0001, ***p<0.001). (C) Silencer activity of the DRL *in vivo*. Top: schematics of the putative silencer placed between the Z48 enhancer and the cardiac actin promoter, in the reporter vector. Intermediate: graphical representation showing EGFP fluorescence intensity signal in midbrain and somites, on the left triggered by FragT14-S and FragT-S-MA, and on the right by Frag1-S and Frag3-S sequences, at 24 hpf; negative control - empty vector; data represented as mean ± SD; n=60 zebrafish embryos/condition from three replicates, Kruskal–Wallis test followed by Dunn’s post-hoc test; ****p<0.0001, ***p<0.001, *p<0.05. Bottom: representative stereomicroscopic images of EGFP expression in zebrafish midbrain and somites driven by FragT14-S, FragT-S-MA, Frag1-S and Frag3-S putative silencer elements; scale bars = 200 µm; arrows indicate EGFP expression in midbrain. Asterisks show area of autofluorescence. See also Figure S5.

These results demonstrate that the DRL region acts as an interacting hub for other CREs, a function that is mostly restricted to the Frag3-AS sequence from the DRL.

### The DRL Functions as a Silencer Outside the *DAB1* Expression Domain

Cis-regulatory elements can have bifunctional activity, driving transcriptional gene activation, as enhancers in certain cell types, and repressing gene expression, functioning as silencers in other cells^64,65^. Consequently, we explored whether the DRL acts as a silencer in non-neural human cells. To achieve this goal, we cloned the non-pathogenic FragT14-S and pathogenic FragT-S-MA DRL-sense sequences upstream of a ubiquitous promoter (TK) in the control of a reporter gene (*luciferase*), in a mammalian expression vector. Of note, FragT14-S showed no ability to drive transcription, as observed in Figure S4A, making it perfect to test silencer potential. To perform the assay, we selected the human HEK293T cells, which express no DAB1 protein (Human Protein Atlas). After transfection, we found that this fragment, either with non-pathogenic (FragT14-S) or pathogenic (FragT-S-MA) sequences, was able to significantly repress TK promoter activity and lower *luciferase* expression compared to the control (p<0.001) (Figure 3B). FragT-S-MA, containing the pathogenic sequence, showed slightly higher *luciferase* expression levels compared with the non-pathogenic sequences FragT14-S and FragT126-S (with 126 repeats).

In light of this result, we wondered if the lack of *DAB1* expression in HEK293T cells is due to DRL active repression. Therefore, we deleted the endogenous DRL from the genome of HEK293T cells (Figures S5A and S5B). In these DRL deleted cells, we observed no significant changes in the expression of *DAB1* or any other genes within its regulatory landscape (Figure S5C). This suggests that the transcription of *DAB1* in HEK293T cells is not solely due to direct repression by the DRL, but instead results from the limited presence of upstream activators of *DAB1*.

We also tested which sub-fragments of the DRL function as silencers. We found that both Frag1-S and Frag3-S were able to repress TK promoter activity close to the background levels (0.4-fold, p<0.001 and 0.2-fold, p<0.0001, compared to the negative control, respectively), with Frag3-S presenting higher potential to repress transcription. Finally, Frag2-S, containing a short non-pathogenic allele (14 repeats), did not silence *luciferase* gene expression in HEK293T cells (Figure 3B).

Next, to understand if the DRL silencer activity is tissue-specific *in vivo*, we cloned FragT14-S, in a *Tol2* transposable element, containing the Z48 midbrain enhancer and a cardiac actin promoter, that drives transcription in embryo somites, upstream of an EGFP reporter. After transgenesis, we observed that FragT14-S decreases the intensity of EGFP expression in the midbrain (Figure 3C), even when harboring the pathogenic repeat (FragT-S-MA), showing that the DRL works as a silencer *in vivo.* Notably, a decrease of EGFP expression in somites was not observed for FragT14-S or FragT-S-MA sequences. Instead, EGFP expression was higher in FragT-S-MA when compared to FragT14-S or the negative control (empty vector; Figure 3C). These results indicate that, in embryo somites where the DRL showed already the ability to drive transcription (Figure S4C), FragT-S-MA was able to drive higher transcription levels than FragT14-S. Moreover, we observed that Frag3-S drives repression in the midbrain, while Frag1-S enhances transcription in somites (Figure 3C); this enhanced function was only clearly detected when the full DRL sequence contained the pathogenic repeat (FragT-S-MA).

Overall, these results show that the DRL functions as a bifunctional regulatory module in brain and muscle, simultaneously enhancing and restricting gene expression in a specific spatial/temporal pattern.

### The DRL is an Enhancer of *DAB1* in human neural stem cells

To investigate further the cis-regulatory role of the DRL, we assessed the transcriptional enhancer ability of this region by dual luciferase reporter assays in cells that express *DAB1*. Thus, we cloned the fragment that alone was unable to drive expression, FragT14-S (Figure S4A), and FragT-S-MA, along with sub-fragments of these sequences (Figure 4A and Figure S5D), into a vector containing the firefly *luciferase* coding sequence under the transcriptional control of a minimal promoter. The enhancer activity of each sequence was assayed in human neural stem cells (hNSC), which showed clear DAB1 expression (Figure S5E). We found that FragT14-S and FragT-S-MA have the ability to activate luciferase expression, compared with the control sequence (Figure 4B; 3- and 7-folds, respectively; p<0.05), showing that the DRL is an enhancer in cells that express *DAB1*. Of note, Frag1-S showed a tendency to activate luciferase expression (4-fold; p>0.05), contrasting with the function of Frag3-S, which was not able to activate luciferase expression (Figure S5F). Finally, when assessing the potential enhancer activity of Frag2-S, both with the pathogenic (Frag2-S-MA) and non-pathogenic repeat (Frag2-S), alone and in combination with Frag1-S or Frag3-S, we found that the differences were not statistically significant compared to the negative control (Figure S5F).

**Figure 4.**
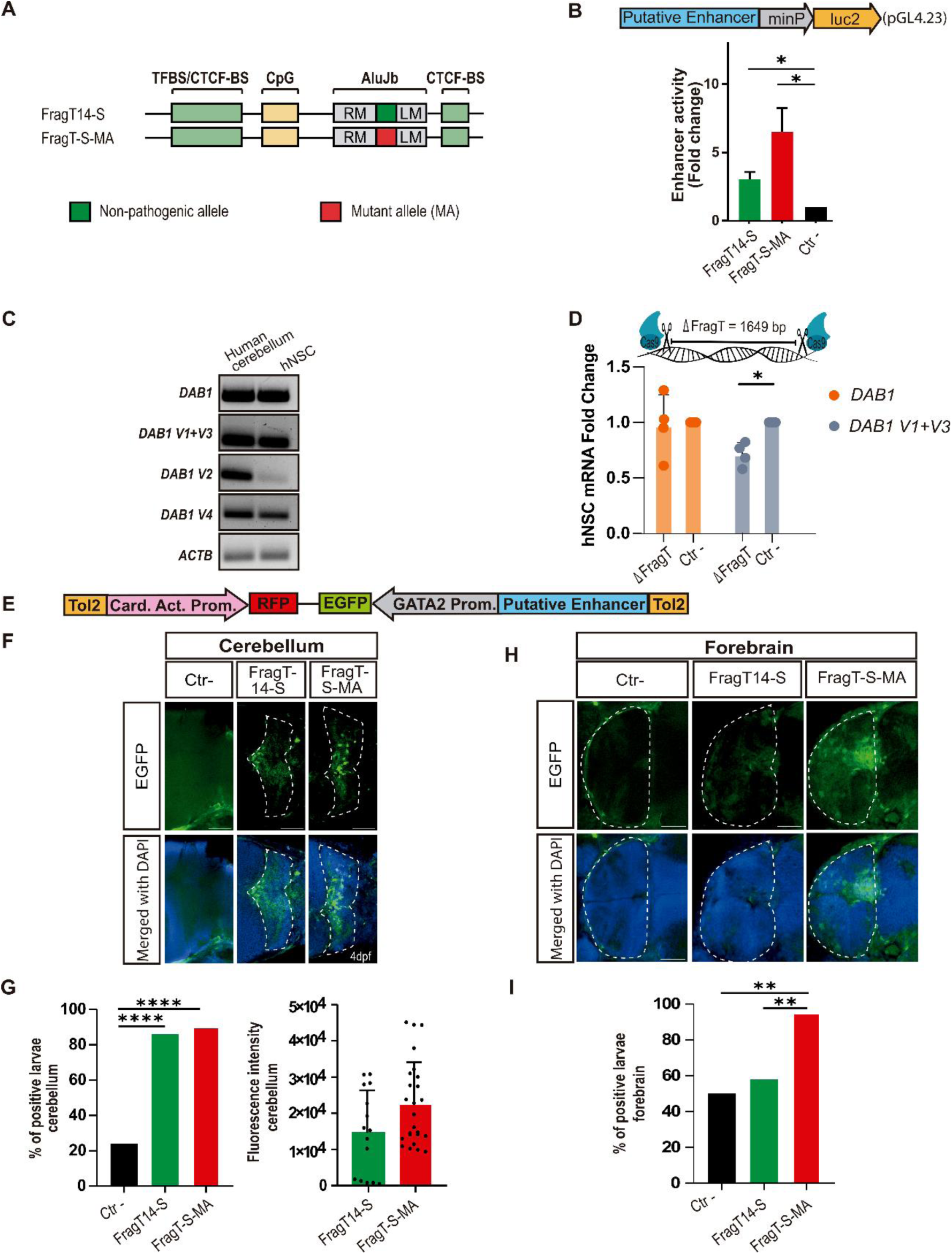
The DRL is Enhancer of *DAB1* in human neural stem cells. (A) Schematics of DRL sequences cloned upstream of a minimal promoter in the control of *luciferase* expression for transcriptional enhancer assay in hNSC. (B) Scheme of reporter vector (top) and graphical representation of assay in hNSC (bottom); enhancer activity is measured by the Luc2/Nluc expression ratios relative to the negative control (ctr-), (data is represented by mean± SD of three biological replicates; independent t-test, * p<0.05); (C) Expression in hNSC cell line, by semi-quantitative RT-PCR, using human cerebellum total RNA as positive control. (D) Real-time quantitative RT-PCR of *DAB1* and *DAB1* V1+V3 variants mRNA from hNSC co-transfected with CRISPR/Cas9 sgRNAs, for DRL deletion (ΔFragT), relative to untransfected cells (ctr-) (four biological replicates; independent t-test, * p<0.05). (E) Schematics of the ZED vector used to *in vivo* validate the enhancer assay by transgenesis in zebrafish embryos; empty vector was used as negative control (ctr-). (F) Representative confocal images of transgenic F0 larvae, 4dpf, showing enhancer activity, in the form of EGFP expression (green), within the cerebellum (dash line area); scale bar=50 μm. (G) Left: percentage of larvae with EGFP positive expression compared with ctr-(29 larvae for FragT14-S; 29 larvae for FragT-S-MA; and 17 larvae for empty ZED vector; Fisher’s exact test, ****p<0.0001). Right: graphic representation of EGFP fluorescence intensity in cerebellum (15 larvae for FragT14-S and 25 for FragT-S-MA). (H) Representative confocal images of transgenic F0 larvae, 4dpf, showing enhancer activity, denoted by EGFP expression, in the forebrain (dash line area), compared with ctr-(scale bar=50 μm). (I) Graphic representation of the percentage of larvae with EGFP expression in zebrafish forebrain compared with ctr-(24 larvae for FragT14-S, 22 larvae for FragT-S-MA and 10 larvae for the empty ZED vector; Fisher’s exact test, **p<0.01). See also Figure S5.

Overall, these results demonstrate that the DRL acts as an enhancer in human DAB1-expressing cells, and that its enhancer activity in neural cells increases when the DRL contains the pathogenic repeat.

To investigate if the DRL sequence is a transcriptional enhancer of *DAB1,* we carried out the deletion of this region by CRISPR/Cas9 in hNSC (Figure S5A). These cells are relevant for this assay because they have expression levels of the *DAB1* variants V1+V3 and V4 comparable to human cerebellum (Figure 4C). After DRL deletion (Figure S5G), we observed that the expression level of *DAB1* variant V1+V3 was significantly decreased compared with non-deleted hNSC (Figure 4D). Other genes in the same genomic proximity, like *MYSM1* and *OMA1*, with expression in hNSC but that do not show clear interactions by 4C-seq with the DRL (Figure 1C), presented levels of expression not reaching statistical significance compared with expression levels in non-deleted cells (Figure S5H).

Overall, these results demonstrate that the DRL is a transcriptional enhancer of *DAB1*, in particular for two transcript variants (V1 and V3) that are highly expressed in human cerebellum and share a promoter that contains the DRL in its regulatory landscape.

### The pathogenic SCA37 DRL Acts as a Hyperactive Brain Enhancer

The most affected brain region in SCA37 is the cerebellum, where *DAB1* is highly expressed from human fetal development to adult life^8^. We demonstrated above that the DRL is an enhancer of cerebellar-specific *DAB1* isoforms, so to determine whether the DRL acts *in vivo* as an enhancer in the cerebellum, we conducted enhancer assays using zebrafish larvae and the zebrafish enhancer detection (ZED) vector^49^. We cloned FragT14-S and FragT-S-MA fragments upstream of a minimal promoter that controls transcription of EGFP (Figure 4E). After genomic integration, we confirmed that both FragT14-S and FragT-S-MA sequences showed consistent EGFP expression in the cerebellum, contrasting with the negative control, demonstrating that the DRL is indeed a cerebellar enhancer (Figure 4F). Importantly, we observed that FragT-S-MA is able to drive higher levels of EGFP intensity than FragT-S (Figure 4G), suggesting that the SCA37 DRL is able to drive higher levels of expression of target genes. Additionally, we found that the FragT-S-MA sequence, but not FragT-S, increased expression of EGFP in other brain regions, such as the forebrain (Figures 4H and 4I), indicating that it acts as a stronger and broader brain enhancer than its non-pathogenic counterpart.

These results demonstrate that the DRL harboring the SCA37 pathogenic repeat has the potential to drive higher levels of expression of *DAB1* in the cerebellum and activate its expression in other regions of the brain, culminating in the *DAB1* expression dysregulation seen in SCA37 brain cells^9^.

### *DAB1* upregulation and dysregulation of reelin pathway in SCA37 neural cells

To investigate neuronal transcriptional deregulation of genes, including *DAB1*, in cells from patients carrying the SCA37 DRL, we differentiated SCA37-derived iPSC in glutamatergic cortical iPSN (Figure 5A) for cap-analysis of gene expression (CAGE). These iPSN showed expression of the neuronal markers TUJ1 and MAP2 (Figure 5B and Figure S6A), confirming their neural identity, as well as DAB1 protein (Figure 5C and Figure S6B). Our analysis detected 3,134 transcripts differentially expressed (DE) between SCA37 iPSN and control iPSN derived from unaffected individuals with p-value <0.05 (Figure 6A). We performed functional annotation analysis for the 457 genes found deregulated with p value<0.05 and -1.5<log_2_(FC)>1.5 in DAVID (Figures 6B, 6C, 6D and 6E). These differentially expressed genes participated mainly in protein-protein interaction and neurotransmission at synapses, according to the TOP10 gene ontology (GO) Reactome pathways (Figure 6B). The TOP10 GO for molecular function indicated they have mostly metal ion and protein binding functions (Figure 6C) and, as expected, they encode components of axons, synapses, dendrites, plasma membrane and nucleus, shown by the GO for cellular components (Figure 6D). Remarkably, these deregulated genes intervene in processes such as nervous system development and axon guidance (Figure 6E). Several of the DE genes belong to the Reelin/DAB1 signaling pathway^66^, including cerebellar-specific *DAB1* variants V1 and V3 (transcribed from the same promoter) (Figure 6F). Some DE genes are associated with other neurodegenerative diseases (Figure 6A), such as *MATR3*^67^ to amyotrophic lateral sclerosis (ALS21), or *SYNE1*^68^ (SCAR8), *TTBK2*^69^ (SCA11) and *PPP2R2B*^70^ (SCA12) to hereditary ataxias.

**Figure 5.**
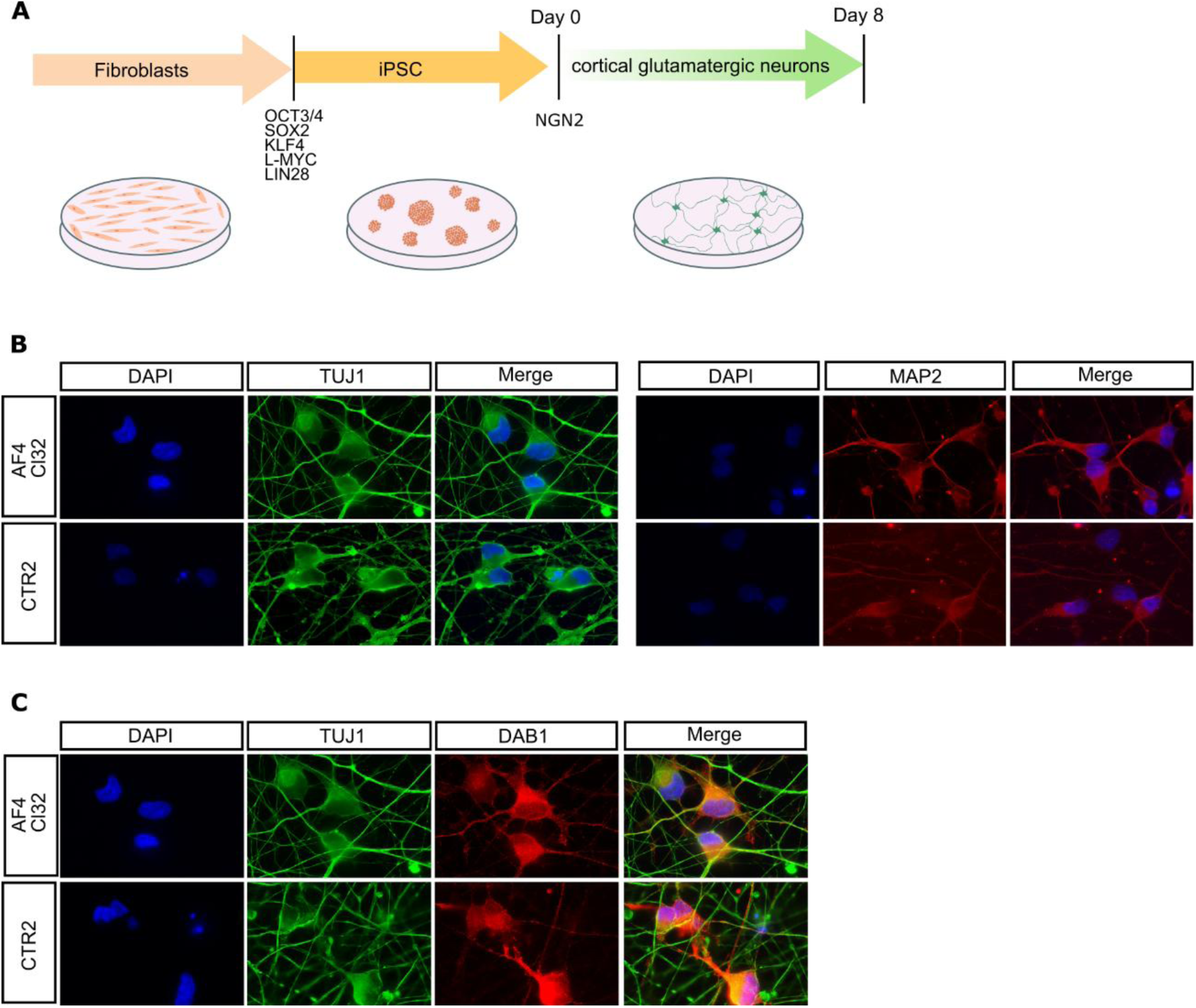
iPSN expressing neuronal-specific markers and DAB1. (A) Schematic representation showing the process used for differentiation in cortical glutamatergic neurons (iPSN) from reprogrammed fibroblasts. Reprogramming of fibroblasts in iPSC was achieved by overexpression of OCT3/4, SOX2, KLF4, L-MYC and LIN28. The iPSN were obtained from iPSC upon NGN2 overexpression (for 8 days) for neuronal differentiation. (B) Expression of neuronal-specific markers, TUJ1 and MAP2, in SCA37 (AF4 Cl32) and unaffected control (CTR2) iPSNs. (C) DAB1 expression in these iPSNs. See also Figure S6.

**Figure 6.**
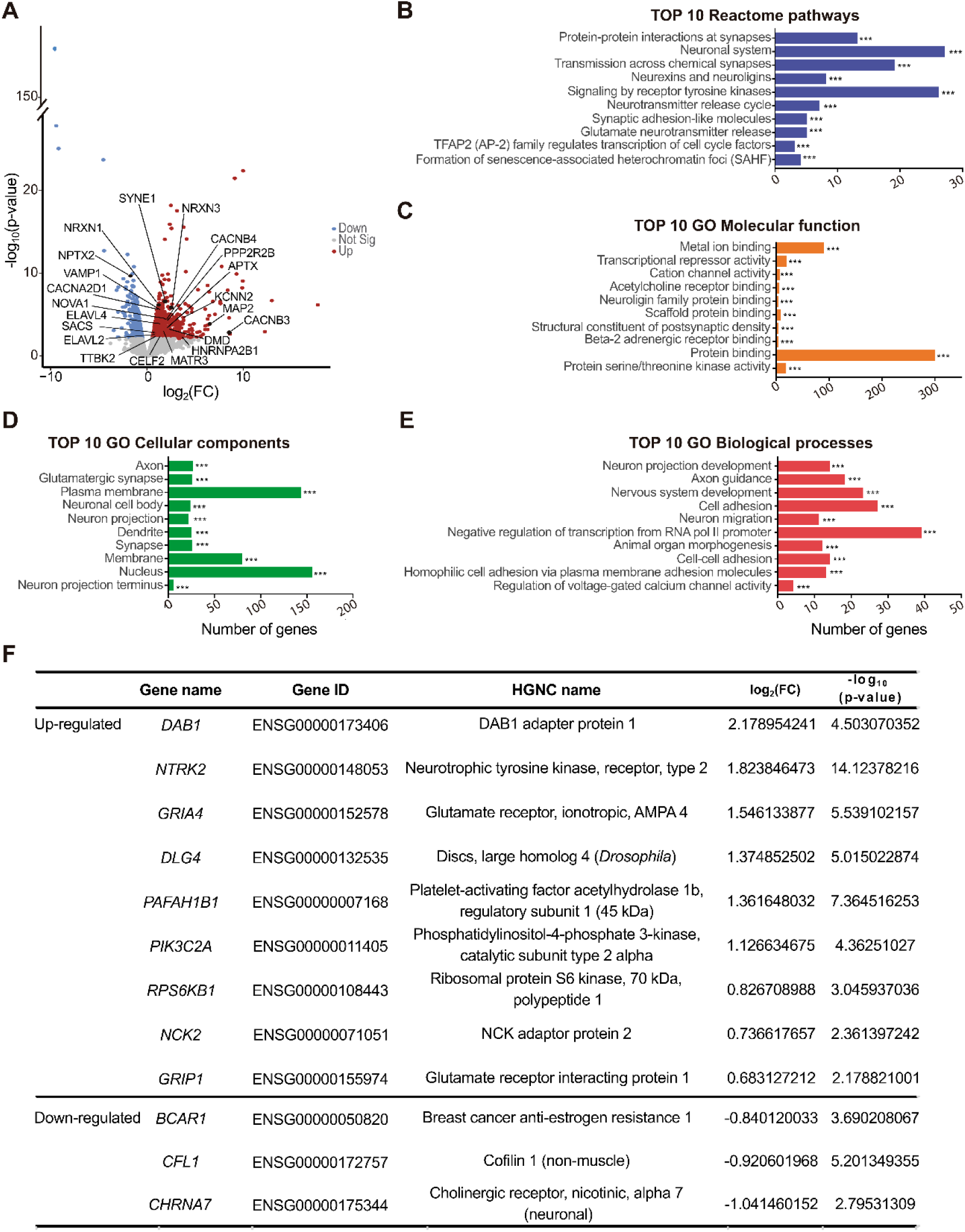
Upregulation of *DAB1* and dysregulation of Reelin pathway in SCA37. (A) Volcano plot showing all deregulated genes found in SCA37-derived iPSN; red-upregulated genes (p<0.05), blue-downregulated genes (p<0.05), grey-no statistical significance. (B-E) DAVID functional annotation of the 457 DE genes in SCA37, with p<0.05, and -1.5<Log_2_(FC)>1.5. (B) TOP 10 reactome pathways (C) TOP 10 categories of gene ontology (GO) for molecular function. (D) TOP10 GO cellular components. (E) TOP10 GO biological processes. (F) Differentially expressed genes from the Reelin/DAB1 pathway. *** p<0.001, p-value of the probability of genes, out of the total genes, annotated to the particular GO term given the proportion of genes in the whole genome that are annotated to that GO term.

These results show that *DAB1* expression is elevated in SCA37 iPSN cells, potentially resulting from the hyperactivity of the DRL enhancer containing the pathogenic SCA37 repeat. Furthermore, this upregulation of *DAB1* likely contributes to the altered expression of genes critical for neuronal function, highlighting a possible cascade effect of ectopic *DAB1* expression on SCA37 early progression.

### DAB1 Overexpression Induces Axonal Defects During Development

Axonal defects are a hallmark of neurodegenerative diseases, namely SCAs^71^. The reelin/Dab1 pathway components are enriched in axonal growth cones of fetal rat brain, where tyrosine phosphorylation of Dab1 is increased compared to other neuronal locations^21^. To investigate if *DAB1* overexpression compromises developmental axonal growth, we microinjected human *DAB1* mRNA into zebrafish embryos and analyzed axonal length in the central and peripheral nervous system. The zebrafish, like the human cerebellum, is composed of an organized three-layer structure with glutamatergic granule cells (GC) innervating GABAergic Purkinje cells (PC) (Figure 7A)^33,72^. After *DAB1* mRNA microinjection, we confirmed translation into DAB1 protein (Figure S7) and analyzed the presence of the longest axons from GC that innervate PC-like cells. We found a significant number of cerebella (identified using a cerebellar GC GFP^+^-expressing zebrafish line^33^) with unilateral or bilateral absence of these axonal tracts compared with control larvae, at 5 dpf (0% for control, 9% for *DAB1* mRNA; p < 0.001; Figures 7B and 7C). To further explore defects in axonal tracts earlier in development, we assessed the length of spinal primary motor neuron (PMN) axons that innervate muscle fibers, at 24 hpf (Figure 7D). We observed that the length of these axonal tracts had extended to a shorter distance in embryos with ectopic expression of DAB1 than in control embryos (40±12 µm for control, 33±10 µm for DAB1; p < 0.05; Figures 7E and 7F).

**Figure 7.**
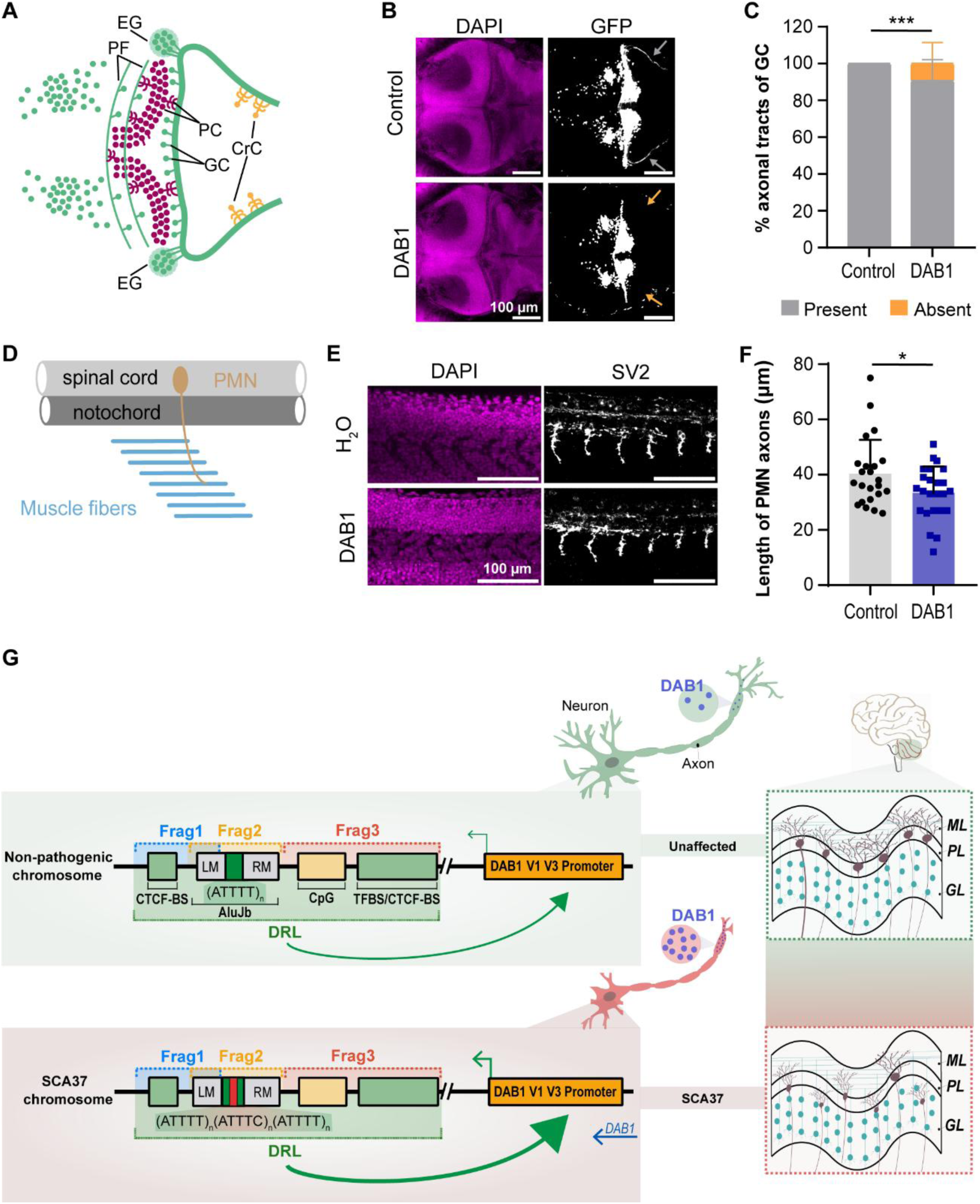
DAB1 overexpression induces axonal defects during development. (A) Schematic representation of cerebellar and cerebellar-like structures in zebrafish. Granule cells (GC) axons, named parallel fibers (PF), innervate Purkinje cells (PC) in the molecular cell layer. Axons from GC in *eminentia granularis* (EG) innervate a type of Purkinje-like neurons called crest cells (CrC). (B) Representative confocal images of zebrafish larvae, from a GFP^+^ cerebellar granule cell line, microinjected with control or human *DAB1* mRNA, at 5 dpf; axonal tracts of GC from EG are indicated with orange (absent) or gray (present) arrows. (C) Percentage of present or absent GC axonal tracts that innervate PC-like neurons (graphic representation of data by mean ± SD of at least 20 embryos/condition/replicate; three independent experiments; ꭕ^2^ test for % of GC axonal tracts, present or absent, ***p<0.001). (D) Schematics of muscle innervation by primary motor neuron (PMN) axon tracts, in zebrafish larvae, at 24 hpf. (E) Representative confocal images (40x objective) of PMN axon tracts from the 6-somites spanning region anterior to the cloaca, from control or *DAB1* mRNA-injected embryos; whole-mounted immunohistochemistry with SV2 antibody, representative z-projection images (maximum intensity). (F) Length of zebrafish PMN axon tracts in control and *DAB1* mRNA microinjected embryos (n=23 embryos for control, n=24 embryos in *DAB1* mRNA condition; four independent experiments; independent t-test, *p<0.05; data represented by mean ± SD. (G) Proposed mechanistic model showing disruption of DRL function by the (ATTTC)_n_, which hyperactivates the neurodevelopmental enhancer, leading to *DAB1* upregulation and DAB1 protein accumulation in axons, likely contributing to SCA37. See also Figure S7.

These results demonstrate that the overexpression of DAB1 during development leads to defects in axonal tracts both in central and peripheral nervous system. Altogether, these results establish a deleterious role for the hyperactivation of the DRL cis-regulatory region in the molecular mechanism leading to SCA37 (Figure 7G).

## Discussion

We report here a primate-specific neurodevelopmental CRE of the human *DAB1* gene, the DRL. The DRL interacts with cerebellar-specific *DAB1* promoters in iPSC and enhances *DAB1* transcriptional expression in hNSC. In SCA37, the pathogenic (ATTTC)_n_ disrupts the DRL, leading to enhancer hyperactivation *in vivo*, and *DAB1* upregulation in SCA37-derived neurons. As DAB1 overexpression during zebrafish neurodevelopment impairs axonal pathfinding, we propose that *DAB1* upregulation contributes to SCA37 pathogenesis.

This work shows the crucial role of Alu STRs in tuning gene expression of proteins functioning in signaling pathways leading, among other processes, to brain development and neurotransmission that ultimately modulate motor and cognitive behaviors. Here, disruption of the Alu STR within the DRL, in SCA37, causes loss of tight *DAB1* expression regulation in neuronal cells, and dysregulation of several other genes in the Reelin/DAB1 signaling pathway. Consistently, DAB1 upregulation has been reported in cerebellar brain tissue from elderly SCA37 individuals^9^, which strongly suggests that this neurodevelopmental enhancer is hyperactive throughout life, in SCA37 subjects.

The disruption of the Alu STR within the DRL shows that SCA37 originates from a complex combination of mechanisms occurring both at DNA and RNA levels. The processes associated with transcribed STR expansions causing neurodegenerative and neuromuscular diseases include mainly the formation of abnormal nuclear RNA aggregates that sequester RNA-binding proteins (RBP) from their cellular location, leading to their loss-of-function^73,74^. We have previously shown that the *DAB1* pathogenic RNA is toxic *in vivo*, causing lethal developmental malformations in zebrafish embryos^8^. We now reveal how disruption of the DRL at the chromatin level leads to *DAB1* upregulation in SCA37 neurons. This disruption provides higher cellular levels of toxic RNA repeat, amplifying this RNA-mediated mechanism. These higher levels of toxic STR-containing RNA could abnormally interact with nuclear RBPs, being more retained in this compartment, modifying cell nucleus architecture and consequently gene expression, like other similar RNAs^75–77^.

The deregulated genes associated with neurodegenerative diseases found in SCA37-derived neurons strongly indicate that the disrupted DRL interferes with their transcriptional regulation. They are involved in many crucial neuronal functions, including axonal guidance. Moreover, recent findings have shown increased complexity in the regulation of gene expression resulting from interaction of RNAs and transcription factors^78^. These deregulated genes associated with other nervous system conditions could explain the clinical overlapping phenotype among SCA37 and other disorders^11^.

DAB1 overexpression in zebrafish embryos showed neurodevelopmental defects in innervation not only in the cerebellum but also in muscle, demonstrating the deleterious effect of upregulated DAB1 protein in the motor system. Moreover, suggesting that the motor neuron pathology seldom observed in affected individuals with the *DAB1* repeat mutation is associated with SCA37^11^. In mammals, *Dab1* encodes a protein highly present in synaptic membrane, presynaptic vesicles and postsynaptic structures^79^ crucial in neurotransmission. During neurodevelopment, Dab1 is located in axonal tracts and growth cones, being required for neurite outgrowth and establishment of neuronal connections^20^. *Dab1* upregulation and protein accumulation in axonal growth cones compromise neurite outgrowth, in Reelin-deficient mice, *reeler*^20^. Together, this strongly suggests that SCA37 is caused by a combination of both DAB1 protein levels dysregulation and a toxic RNA-mediated mechanism.

The neurodevelopmental phenotype associated with *DAB1* upregulation in this study supports the idea of an active role of developmental alterations in the mechanism leading to SCA37. This idea is further strengthened by the very early suggestive evidence of embryonic and molecular abnormalities in human fetal carriers of the neurodegenerative Huntington’s disease STR expansion^80^.

Insertions of (ATTTC)_n_ in antisense Alu elements have also arisen in other genes causing neurological diseases with associated signs and symptoms of cerebellar pathology^5,6^, suggesting a shared pathological mechanism for these conditions. Besides, many nucleotide repeat expansions have originated in Alu elements^5,6^. Moreover, as multiple Alu elements in the human genome carry STRs in their A-rich regions^81^, they could have comparable regulatory roles in other cells and organs that when disrupted might confer risk, as in cancer^82^, or cause diseases such that involving metabolic feedback regulation^83^. We anticipate the epigenomic technologies in use will raise the pace of discovery of regulatory Alu STR regions and related complex mechanisms, improving prevention and providing therapeutic targets.

## Acknowledgements

We are grateful to Drs. Paulo Pereira and João M. Cabral for comments on the manuscript. We are thankful to Dr. Ana Valentim for advice with statistical analysis. We thank Joana Marques and Isabel Guedes for the maintenance and assistance with zebrafish work; Tiago Ribeiro, Marta Duque and Dr. Rita Barbosa Matos for the helpful support with protein detection, imaging processing and karyotype protocols. We are grateful to the i3S Information Systems and Technology Unit for helpful assistance and for providing access to the *totoro* HPC facility. We acknowledge NINDS for providing iPSC control samples. We also acknowledge Prof. Koichi Kawakami from the National Institute of Genetics, and the National BioResouce Project from the Ministry of Education, Culture, Sports, Science and Technology of Japan, as well as Prof. Masahiko Hibi from the Nagoya University, Japan, for providing the zebrafish lines gSA2AzGFF152B and UAS;GFP.

This work was financed by National Funds through FCT—Fundação para a Ciência e a Tecnologia, I.P., under the project UIDB/04293/2020 and by Fundo Europeu de Desenvolvimento Regional (FEDER), through the COMPETE 2020 Operational Program for Competitiveness and Internationalization (POCI) of Portugal 2020, FCT and Ministério da Ciência, Tecnologia e Ensino Superior (Portugal), in the framework of the project POCI-01-0145-FEDER-029255 (PTDC/MED-GEN/29255/2017) to I.S. and J.B. A.F.C and AS.F. are recipients of FCT PhD scholarships (2020.00528.BD and 2021.05757.BD). This work has also received funding from the European Union’s Horizon 2020 research and innovation programme under grant agreement No 952334; and to the Scientific Platform Advanced Light Microscopy, as member of the national infrastructure PPBI-Portuguese Platform of BioImaging (supported by POCI-01-0145-FEDER-022122). The following funding supported also this work: European Research Council (ERC) under the European Union’s Horizon 2020 research and innovation program (ERC-2015-StG-680156-ZPR); “La Caixa” Foundation (under the grant agreement HR21-01212); and Fundação para a Ciência e a Tecnologia (FCT) in the framework of the project PTDC/BIA-MOL/3834/2021 to JB. FCT grant CEECIND/03482/2018 supported JB.

## Author contributions

J.R.L conceived of this project with I.S and J.B., led the development and optimization of cellular experiments, performed experiments, analyzed and interpreted data, wrote the paper and produced the related figures. A.F.C. performed zebrafish experiments, analyzed and interpreted data, wrote the paper and generated the related figures. A.S.F. performed iPSC analysis, HEK293T editing and analysis, contributed to the writing of these sections, model/schematic illustrations and provided comments and edits on the entire manuscript. A.E. performed zebrafish enhancer assay in vivo, contributed to the writing, model schematic/illustrations and provided comments on the manuscript. A.D. performed iPSC and iPSN analysis and provided comments on the manuscript. M.G. performed 4C-seq data analysis and provided comments on the manuscript. H.M. performed zebrafish experiments, analyzed and interpreted data and reviewed the manuscript. C.R. performed HEK293T luciferase assays, analyzed and interpreted data, wrote the paper, produced the related figures and reviewed the manuscript. P.S., M.A. and M.S provided guidance and support in imaging analysis and created new methods for data analysis and visualization, wrote software, and reviewed and edited the manuscript. S.D. performed karyotype analysis and reviewed and edited the manuscript. P.R. conceived and led the development, optimization and analysis of CAGE experiments, provided comments and reviewed and edited the manuscript. P.H. conceived and led the generation and validation of iPSC and iPSN experiments, provided comments and reviewed and edited the manuscript. J.B. conceived of this project with I.S. and J.R.L. and oversaw all the analysis and experiments carried out; wrote the paper. I.S. conceived of this project with J.B. and J.R.L. and oversaw all the analysis and experiments performed; wrote the manuscript.

## Declaration of AI and AI assisted technologies

No AI technologies were used.

