## Supplementary Figures and Tables for "The Insertion of an ATTTC Repeat in an Alu Element Hyperactivates a Primate-Specific Neurodevelopmental Enhancer in Spinocerebellar Ataxia Type 37"

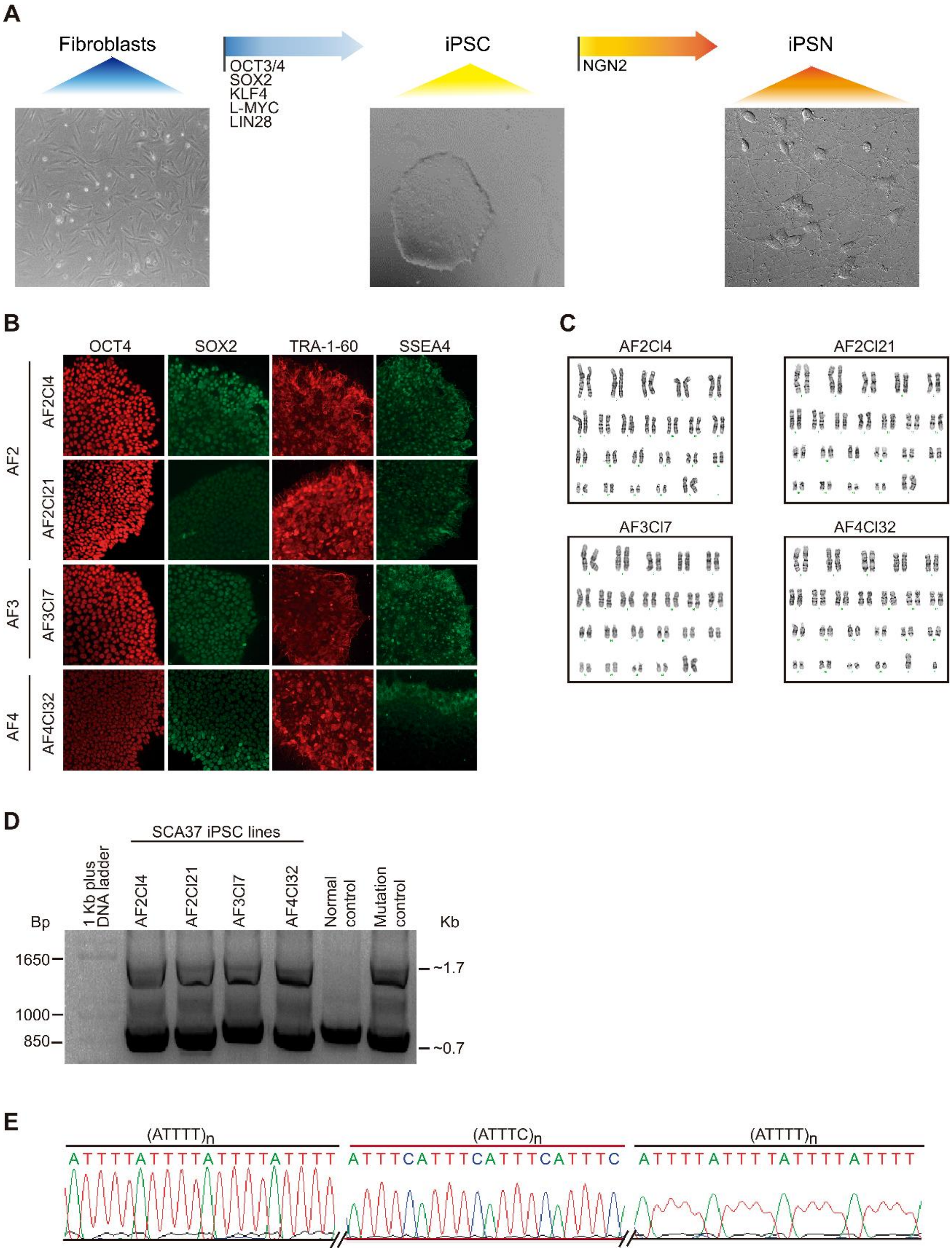


**Figure S1. Generation and characterization of SCA37 iPSC lines**. Related to **Figure 1.** (A) Shown are brightfield microscopy representative images of SCA37-derived fibroblasts before and upon reprogramming in iPSC, by transfection with OCT3/4, SOX2,

KLF4, L-myc and LIN28 reprogramming factors. It also depicts a representative brightfield microscopy image of neurons obtained from iPSC differentiation upon overexpression of NGN2. (B) Expression of pluripotency markers in four iPSC-patient derived SCA37 lines detected by immunofluorescence. (C) Karyotype representative images of the four SCA37 iPSC lines, showing normal chromosome pairs with no aberrant chromosome rearrangements. (D) iPSC genotyping of the four SCA37-iPSCs lines with PCR amplification of unaffected and mutant pentanucleotide alleles, in SCA37. (E) Sanger sequencing of a mutant allele amplified in a representative SCA37 iPSC line, showing the $(ATTTC)_n$ insertion.

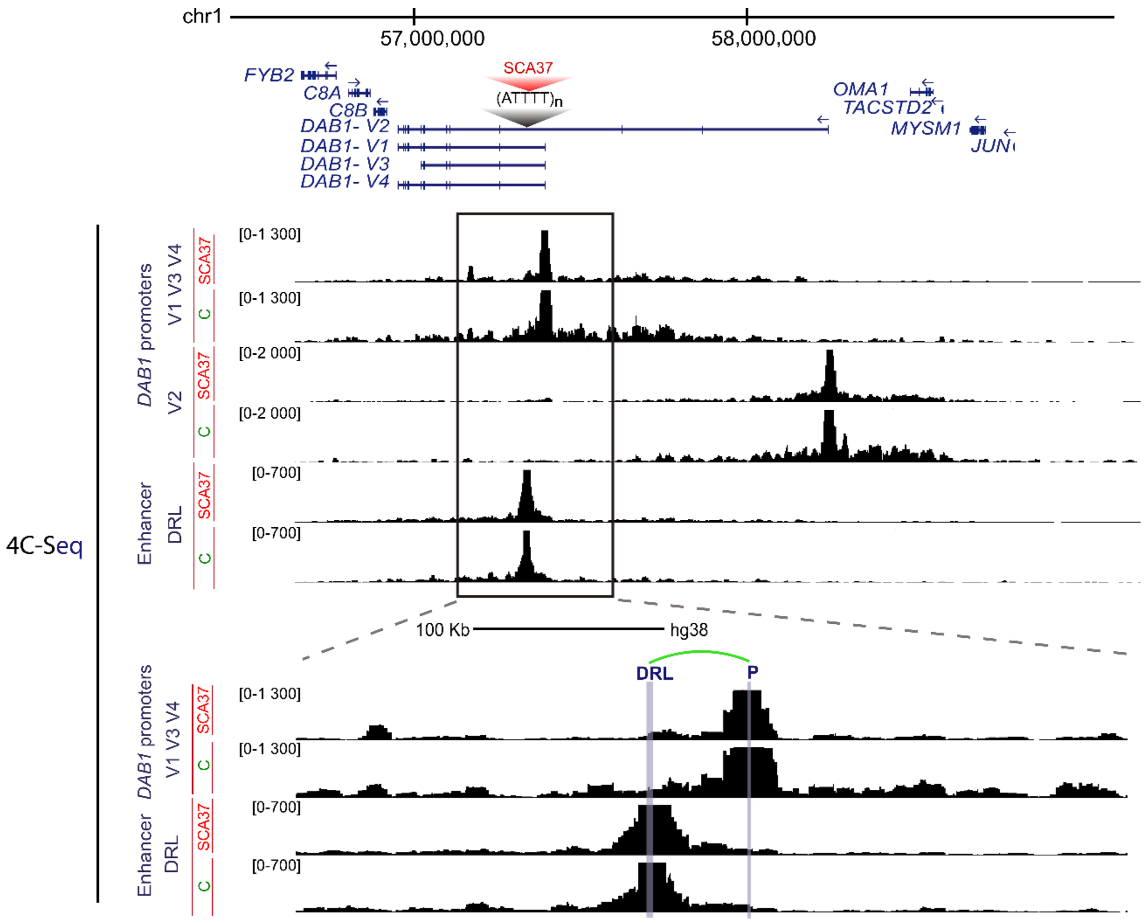


**Figure S2**. **4C-seq of additional SCA37 iPSC and unaffected control lines.** Related to **Figure 1C**. 4C-seq interaction profiles of additional iPSCs-derived from SCA37 and unaffected subjects, with viewpoints in the promoters of *DAB1* V1, V3 and V4 transcript variants and that of V2 *DAB1* transcript, as well as in the DRL region. C-control, P-promoter.

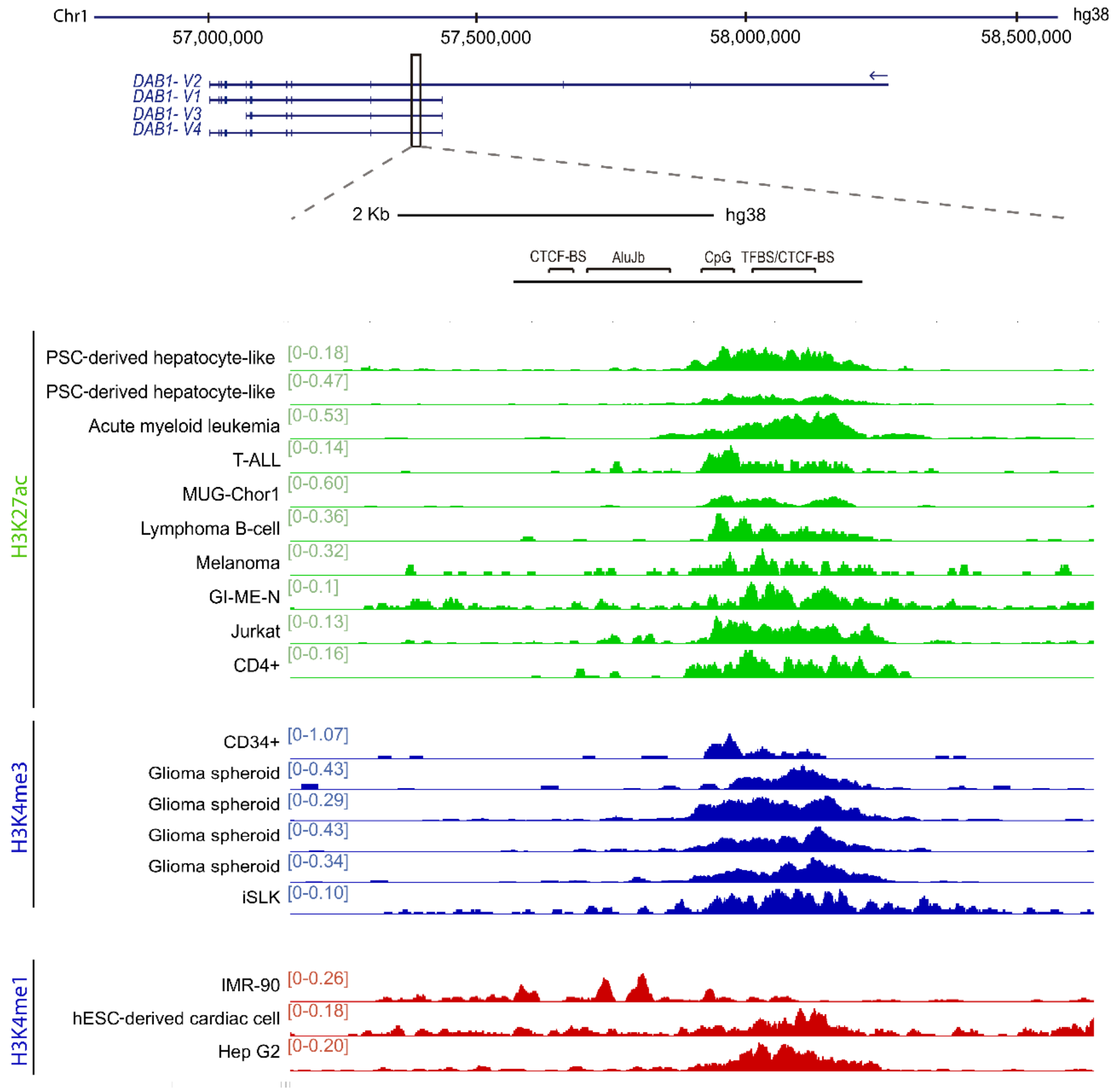


**Figure S3. ChIP-seq histone marks at the DRL genomic region**. Related to **Figure 1D**. Additional human cells with positive ChIP-seq signal for H3K27ac, H3K4me3 and H3K4me1 marks, at the DRL region, from ChIP-Atlas.

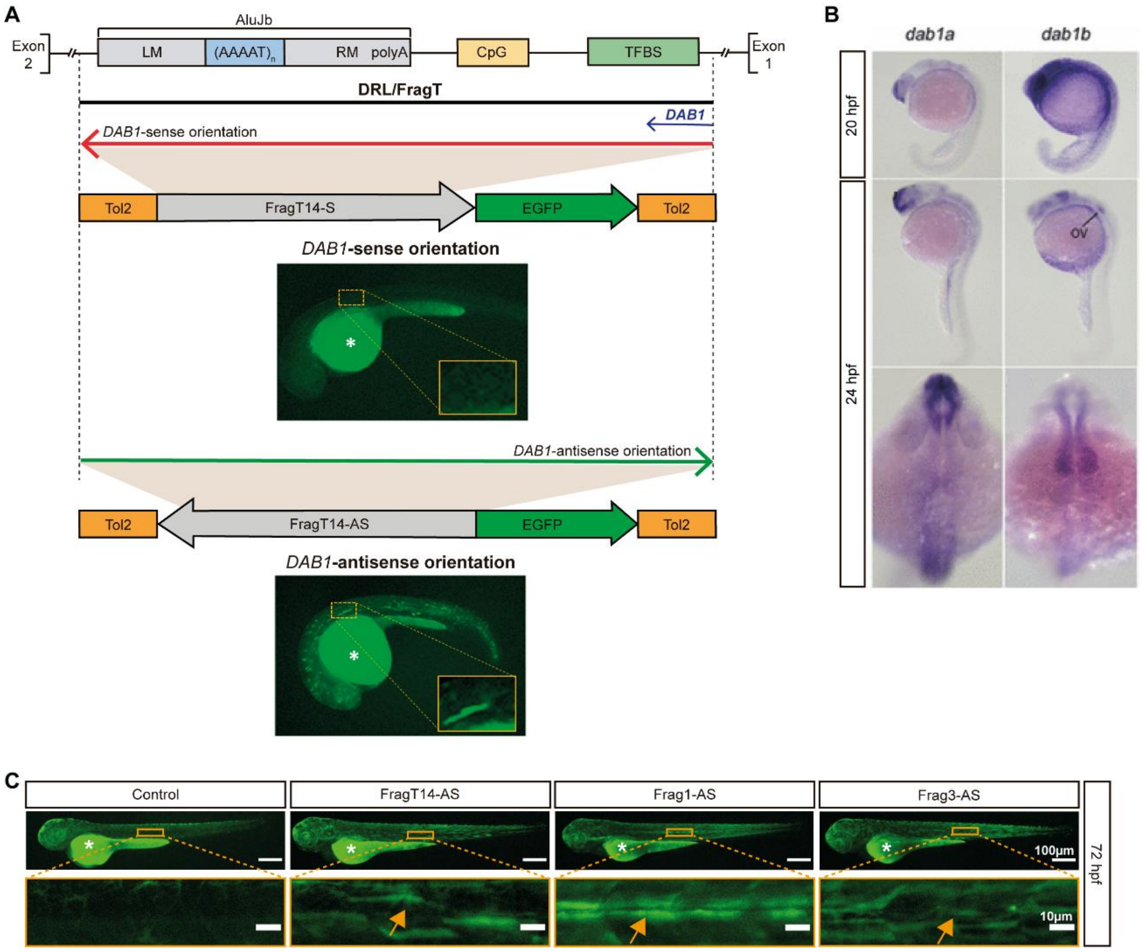


**Figure S4. Neurodevelopmental enhancer activity of the DRL region.** Related to **Figure 2.** (A) Schematic representation of the DRL region (top); no EGPP expression denoting *in vivo* regulatory activity for FragT14-S/DRL region, in zebrafish, at 24hpf (upper panel), in contrast to FraT14-AS, in the *DAB1*-antisense orientation (lower panel). (B) Expression pattern of zebrafish *Dab1* homologues, *dab1a* and *dab1b* adapted from Imai and colleagues[62]. (C) FragT14-AS and Frag3-AS regulatory elements drive expression to muscle fibers (orange arrow) and Frag1-AS triggers expression in the horizontal myoseptum (orange arrow), in zebrafish embryos, at 72 hpf; scale bars =100 µm; zoom-in scale bars: 10 µm. Asterisks indicate autofluorescence.

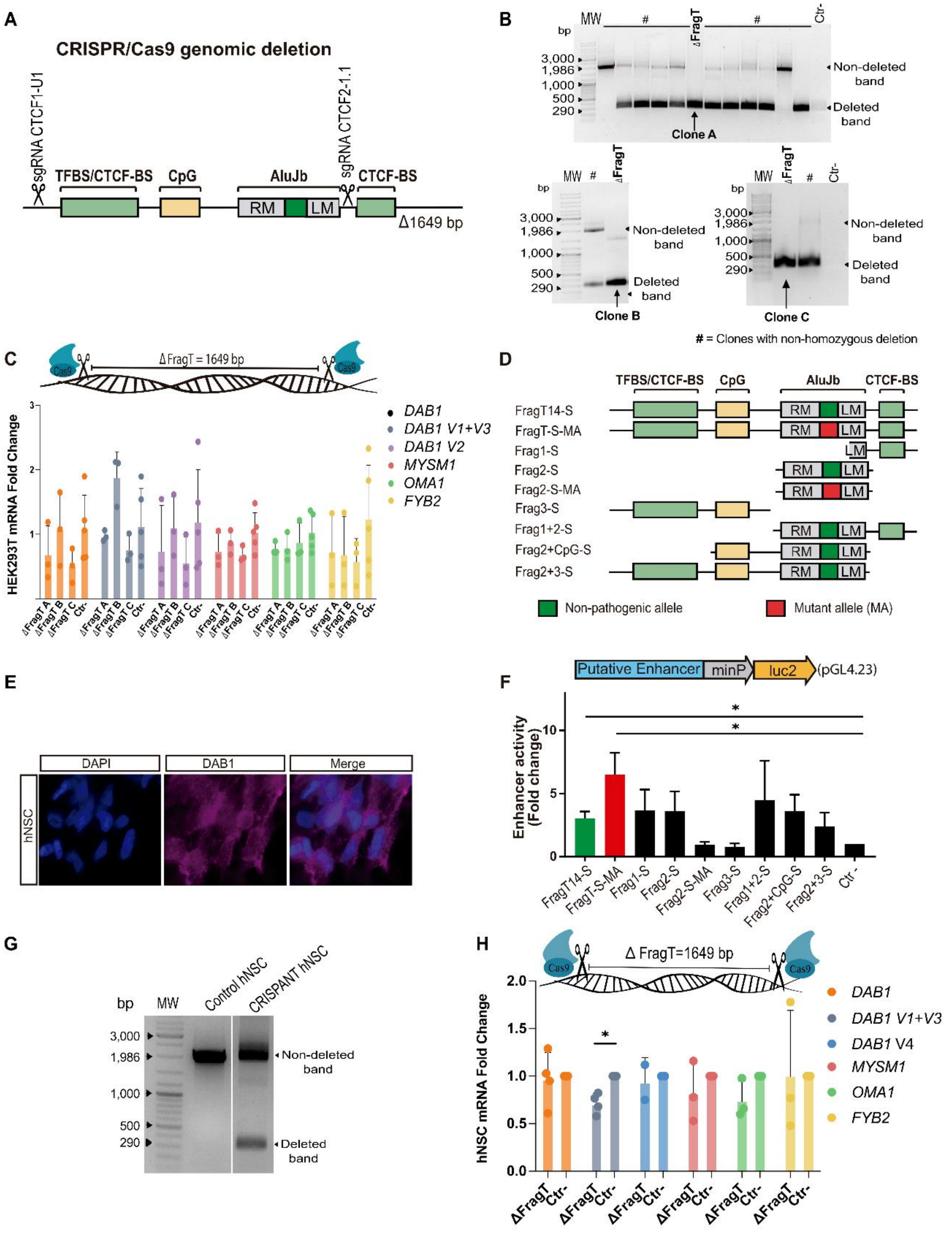


**Figure S5. Genomic deletion of the DRL region in hNSC and HEK293T.** Related to **Figures 3B, 4B and 4D.** (A) Schematic representation of the sgRNA genomic targets used to delete the DRL region, by CRISPR/Cas9, from the genome of hNSC and HEK293T cell lines. (B) PCR analysis of the DRL sequence for validation of the DRL deletion in HEK293T cell clones co-transfected with sgRNAs, arrows indicate the selected clones with the homozygous DRL deletion (ΔFragT). (C) Expression levels assessed by real-time quantitative RT-PCR of *DAB1* and other genes in its landscape, in HEK293T selected clones homozygous for the DRL deletion, relative to control HEK293T cells (ctr-). (D) Schematics of DRL sequences cloned upstream of a minimal promoter in the control of luciferase expression for transcriptional enhancer assay in hNSC. (E) Immunostaining with DAB1 antibody in hNSC, shown by immunofluorescence of DAB1 protein expression (magenta). (F) Scheme of reporter vector (top) and graphical representation of enhancer assay in hNSC (bottom); enhancer activity measured by the Luc2/Nluc expression ratios relative to the negative control (ctr); data is represented by mean ± SD of three biological replicates; independent t-test, *$p<0.05$. (G) PCR amplification of the DRL sequence for validation of the DRL deletion in hNSC co-transfected with sgRNAs; control hNSC were used as a negative control. (H) Expression levels assessed by real-time quantitative RT-PCR of *DAB1* and genes in its landscape, in hNSC co-transfected with CRISPR/Cas9 sgRNAs (ΔFragT) relative to untransfected cells (ctr-), data from four replicates; independent t-test, *$p<0.05$.

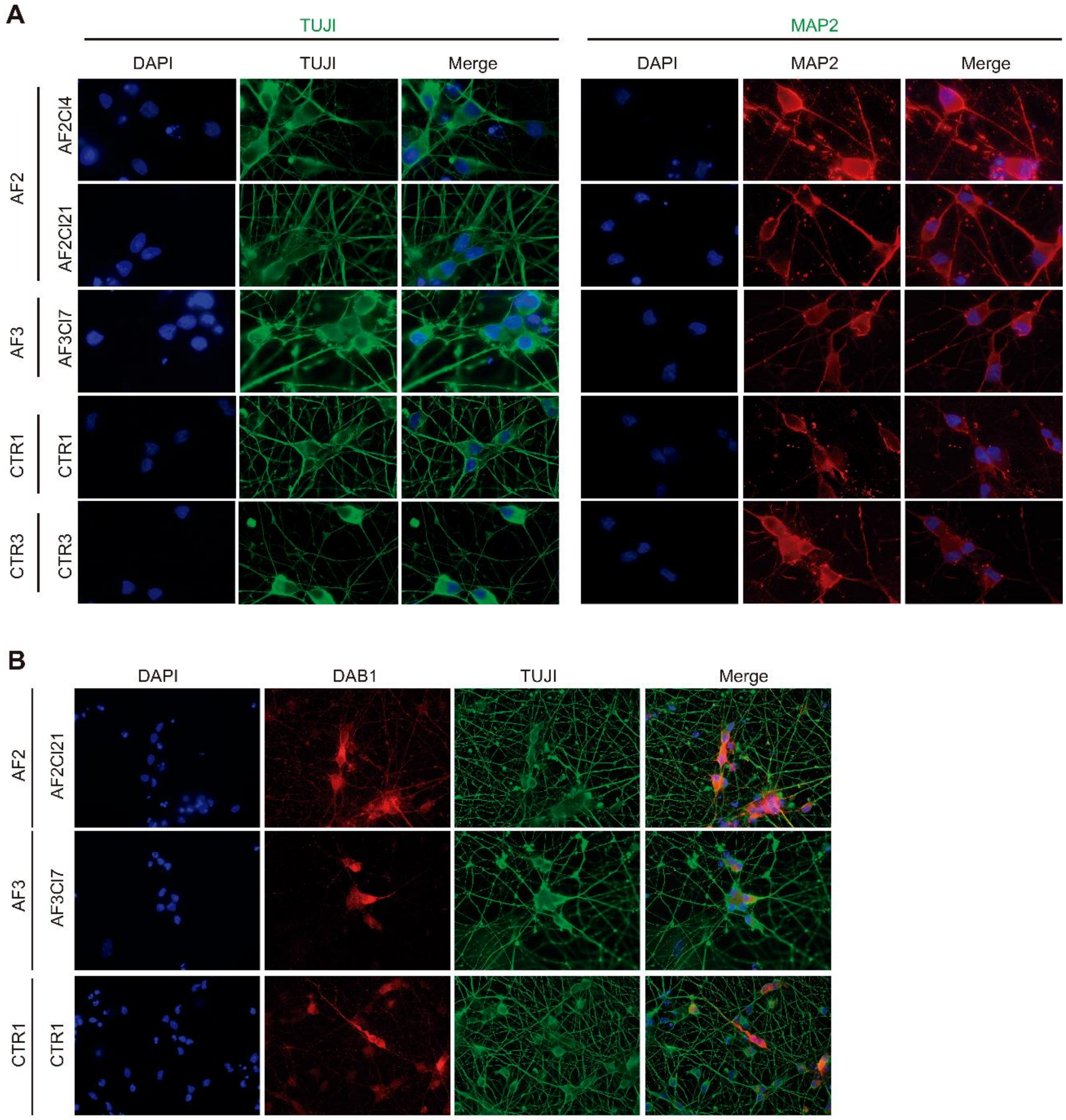


**Figure S6. Three additional iPSN with neuronal-specific markers and DAB1 protein expression**. Related to **Figure 5**. (A) Expression of neuronal markers TUJI and MAP2 in the three additional iPSN differentiated from SCA37 iPSC lines. (B) Detection of DAB1 protein expression in these three additional SCA37 iPSN, by immunofluorescence.

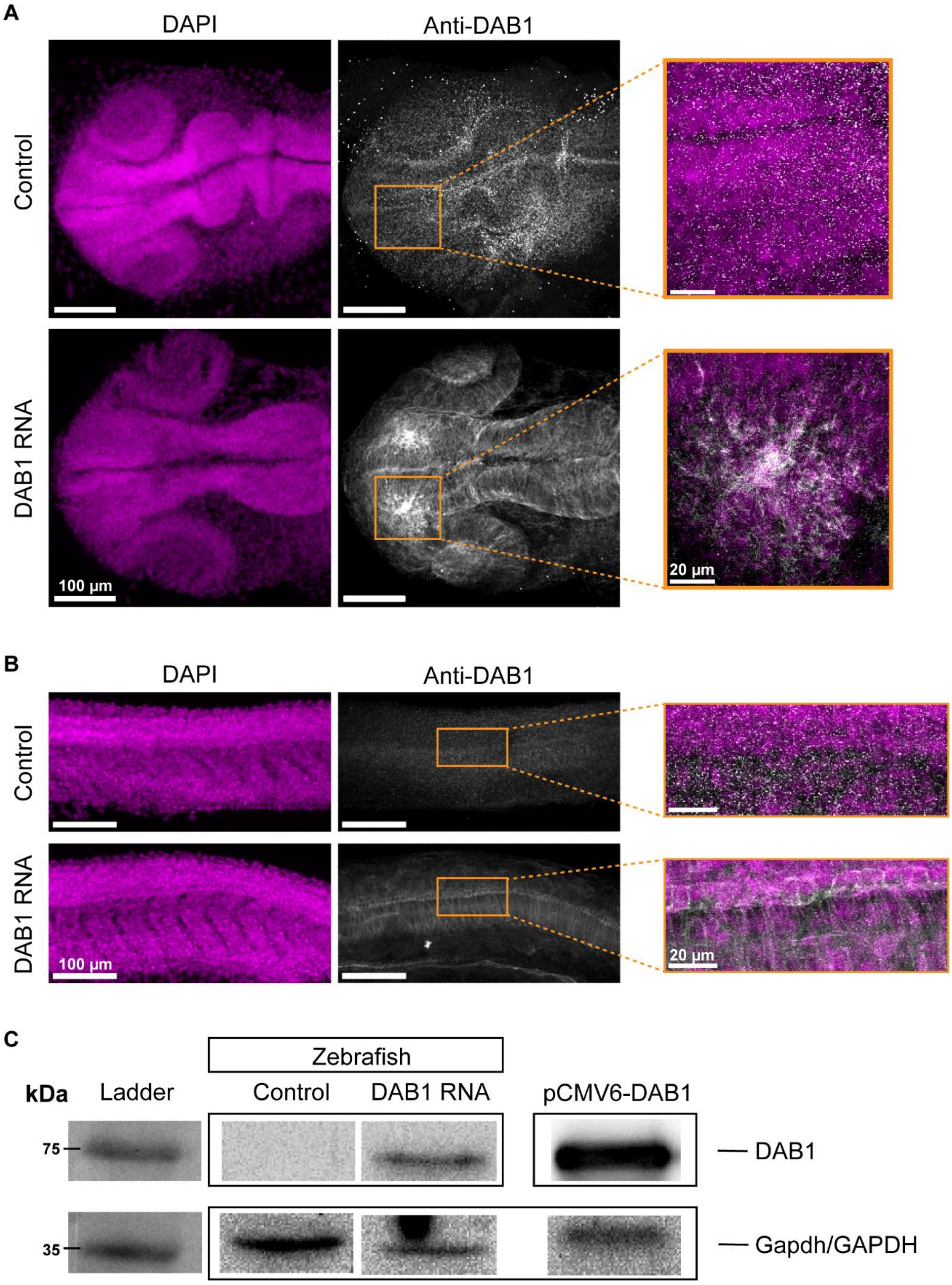


**Figure S7. Translation of human *DAB1* mRNA in microinjected zebrafish embryos.** Related to **Figure 7**. (A and B) Human DAB1 protein expression in zebrafish head (A) and tail (B) after microinjection of control and human *DAB1* mRNA assessed by whole-mount immunohistochemistry with DAB1 antibody, at 24 hpf; representative zprojection images (maximum intensity). (C) Human DAB1 protein expression in zebrafish assessed using anti-DAB1 antibody, by western blot. Protein lysate from HEK293T cells transfected with the human *DAB1* cDNA plasmid pCMV6-DAB1 was used as a positive control. Blot stripped and re-probed with anti-GAPDH antibody for protein loading control. Crop areas relative to different membrane exposure times (DAB1: 5s for zebrafish lysates and 1s for HEK293T cells lysate; GAPDH: 20 min).

**Supplementary Table S1.** Details of the subject iPSC lines derived from SCA37 subjects and controls

| Subject Cell line | Age at Sampling (yrs) | Gender | Phenotype |
|---|---|---|---|
| ND41866 | 64 | Male | Unaffected |
| GM23280 | 36 | Female | Unaffected |
| NAS6 | Unknown | Unknown | Unaffected |
| AF2Cl4 | 40-50 | Figure S1 | SCA37 |
| AF2Cl21 | 40-50 | Figure S1 | SCA37 |
| AF3Cl7 | 50-60 | Figure S1 | SCA37 |
| AF4Cl32 | 40-50 | Figure S1 | SCA37 |

**Supplementary Table S2.** Oligonucleotides for 4C-PCR, cloning, qRT-PCR, genotyping and sgRNA generation.

| Samples | Oligonucleotide ID | Sequence (5´-3') |
|---|---|---|
| **4C PCR** | | |
| SCA37 AF2 and AF4 | p5-V2NR-AF-N503 | AATGATACGGCGACCACCGAGATCTACACTATCCTCTTCGTCGGCAGCGTCAGATGTGTATAAGAGACAGCCTTCTTGGGGTGTTCAC |
| SCA37 AF2 and AF4 | p7-V2R-AF-N705 | CAAGCAGAAGACGGCATACGAGATAGGAGTCCGTCTCGTGGGCTCGGAGATGTGTATAAGAGACAGCTTTGAAAAATGAGATGATTGATC |
| Control ND41866 and GM23280 | p5-V2NR-ND-N505 | AATGATACGGCGACCACCGAGATCTACACGTAAGGAGTCGTCGGCAGCGTCAGATGTGTATAAGAGACAGCCTTCTTGGGGTGTTCAC |
| Control ND41866 and GM23280 | p7-V2R-ND-N701 | CAAGCAGAAGACGGCATACGAGATTCGCCTTAGTCTCGTGGGCTCGGAGATGTGTATAAGAGACAGCTTTGAAAAATGAGATGATTGATC |
| SCA37 AF2 and AF4 | p5-V123R-AF-N502 | AATGATACGGCGACCACCGAGATCTACACCTCTCTATTCGTCGGCAGCGTCAGATGTGTATAAGAGACAGGAACAGAGTCCACAGAGGATC |
| SCA37 AF2 and AF4 | p7-V123NR-AF-N706 | CAAGCAGAAGACGGCATACGAGATCATGCCTAGTCTCGTGGGCTCGGAGATGTGTATAAGAGACAGGCCACTGGGTGAGTGTAC |
| Control ND41866 and GM23280 | p5-V123R-ND-N501 | AATGATACGGCGACCACCGAGATCTACACTAGATCGCTCGTCGGCAGCGTCAGATGTGTATAAGAGACAGGAACAGAGTCCACAGAGGATC |
| Control ND41866 and GM23280 | p7-V123NR-ND-N702 | CAAGCAGAAGACGGCATACGAGATCTAGTACGGTCTCGTGGGCTCGGAGATGTGTATAAGAGACAGGCCACTGGGTGAGTGTAC |
| SCA37 AF2 and AF4 | p5-RepeatNR-AF-N504 | AATGATACGGCGACCACCGAGATCTACACAGAGTAGATCGTCGGCAGCGTCAGATGTGTATAAGAGACAGGAATTATACCTACAAAACTTAACTC |
| SCA37 AF2 and AF4 | p7-RepeatR-AF-N703 | CAAGCAGAAGACGGCATACGAGATTTCTGCCTGTCTCGTGGGCTCGGAGATGTGTATAAGAGACAGGGATGTTAGGGTCAGATC |
| Control ND41866 and GM23280 | p5-RepeatNR-ND-N506 | AATGATACGGCGACCACCGAGATCTACACACTGCATATCGTCGGCAGCGTCAGATGTGTATAAGAGACAGGAATTATACCTACAAAACTTAACTC |
| Control ND41866 and GM23280 | p7-RepeatR-ND-N704 | CAAGCAGAAGACGGCATACGAGATGCTCAGGAGTCTCGTGGGCTCGGAGATGTGTATAAGAGACAGGGATGTTAGGGTCAGATC |
| **Cloning of FragT and subfragments in pMCS or Topo8** | | |
| FragT SCA37 allele | Alu CTCF 2F EcoRI | CCGGCGAATTCTTGTTTCACACATGGCAGTGG |
| | Alu CTCF 2R XmiI | GCACAGTCGACCTGAGCTTGGGGCTTTTATGG |
| | Alu CTCF 2F XmiI | GCATAGTCGACTTGTTTCACACATGGCAGTGG |
| | Alu CTCF 2R EcoRI | GCACAGAATTCCTGAGCTTGGGGCTTTTATGG |

| | | |
|---|---|---|
| FragT Normal allele | Alu CTCF F | TTTCACACATGGCAGTGGCTTA |
| | Alu CTCF R | GGCTGAGCTTGGGGCTTTTA |
| Frag1 | Alu Frag1 F | CACACATGGCAGTGGCTTAC |
| | Alu Frag1 R1 | CTCGAACTCCTGGGCTAAAG |
| Frag2 Normal allele | Alu 24R* | TGAAACTGAGGCTCAAAATGA |
| | Alu 24F* | ATTTGCCCTTTGCTGATTGA |
| Frag2 SCA37 allele | Alu24F_XmiI | GTAGTCGACATTTGCCCTTTGCTGATTGA |
| | Alu24R_EcoRI | GCAGAATTCTGAAACTGAGGCTCAAAATGA |
| Frag3 | Alu Frag2 F | AAAAAGTAAGAGTTCGGAAGTCTG |
| | Alu Frag2 R | ATGGGCCTCAGAGAGAAGTG |
| Frag1+Frag2 | Alu Frag1 F | CACACATGGCAGTGGCTTAC |
| | Alu 24F* | ATTTGCCCTTTGCTGATTGA |
| Frag2+Frag3 | Alu 24R* | TGAAACTGAGGCTCAAAATGA |
| | Frag2 R | ATGGGCCTCAGAGAGAAGTG |
| Frag2+CpG | Alu 24R* | TGAAACTGAGGCTCAAAATGA |
| | CpG Alu 24F | CAGAAGGGACAGTGCTCC |
| **Cloning of TK promoter in pGL4.23-GW** | | |
| TK promoter | TK promoter F XhoI | GTCATCTCGAGAAATGAGTCTTCGGACCTCG |
| | TK promoter R BglII | GCACTAGATCTTTAAGCGGGTCGCTGCAG |
| **qRT-PCR** | | |
| DAB1 all mRNA variants | DAB1_qPCR_v1234_F | TAAAGGTGAAGGGGTCCGGT |
| | DAB1_qPCR_v1234_R | GTTCTCCTTTGGAACGAGCG |
| DAB1 V1 and V3 | RT_15ex1_2_F | AGGATGCTCTGGGCTAGGC |
| | RT_15ex1_2_R | CCAAGAGAAAGACTCCTCCCTTCA |
| DAB1 V2 | RT_17ex_ex3_4_F | CAGCAAGACATGGTGGTGAG |
| | RT_17ex_ex3_4_R | AGAAAGACTCCTCCCTTCAGAAAT |
| DAB1 V4 | UC0009 F1 | CTGTGGTTCGGCTCATAAA |
| | UC0009 R1 | GCTGCACATTTCATTCACTC |
| DAB1 TAD genes | C8AqPCR_F | CTGGATCCATGGGTGGCATT |
| | C8AqPCR_R | CCCAAGGAGCCTCCAAAACA |

| | | |
|---|---|---|
| | FYB2qPCR_F | ACTCTGCCTTGGAGGAAAGC |
| | FYB2qPCR_R | TTGGAGAAACACCTGCTGGG |
| | MYSM1_qPCR_F | GCTGTTTGAACAAGGGCTGG |
| | MYSM1qPCR_R | TCCAGACCGCATTTGACCTT |
| | OMA1qPCR_F | GCCAACCCAAGATGCCAGAA |
| | OMA1qPCR_R | GGTCTGGATTAGACAGTGGTGG |
| | TACSTD2qPCR_F | CTCCAAGTGTCTGCTGCTCA |
| | TACSTD2qPCR_R | GTCGTAGAGGCCATCGTTGT |
| Reference genes | CAPN10 F | CATTCACAGCCAGGAGATGCT |
| | CAPN10 R | CTGTTAGGTTTTCATCACTGCCAT |
| | GNA13 F2 | TCTCCAAGTTTATCCAAGTTATCCAG |
| | GNA13 R2 | TATGACCGGCGTCGAGAAT |
| **Genotyping HEK293T clones for FragT deletion** | | |
| | Alu CTCF 2F | CCGGCGAATTCTTGTTTCACACATGGCAGTGG |
| | Alu CTCF 2R | ATAATGCGGCCGCTGAGCTTGGGGCTTTTATGG |
| **Oligonucleotides for sgRNA generation** | | |
| | Sg_CTCF1_U1F | CACCTTGGCACCCACTTCTCTCTG |
| | Sg_CTCF1_U1R | AAACCAGAGAGAAGTGGGTGCCAA |
| | Sg_CTCF2_1.1F | CACCAAACTCTGCAGCCAGTAGCA |
| | Sg_CTCF2_1.1R | GTGCTGCTACTGCGTGCAGAGTTT |

* Seixas et al, 2017

**Supplementary Table S3.** Sequences of sgRNAs used to delete the FragT/DLR in HEK293T and hNSC cells.

| sgRNA ID | Sequence (5'-3') | PAM sequence | MIT Guide specificity score | Doench score |
|---|---|---|---|---|
| Sg CTCF1_U1 | TTGGCACCCACTTCTCTCTG | AGG | 55 | 62 |
| Sg CTCF2_1.1 | AAACTCTGCAGCCAGTAGCA | TGG | 70 | 59 |
